# Cell type specific control of basolateral amygdala plasticity via entorhinal cortex-driven feedforward inhibition

**DOI:** 10.1101/348524

**Authors:** E. Mae Guthman, Joshua D. Garcia, Ming Ma, Philip Chu, Serapio M. Baca, Katharine R. Smith, Diego Restrepo, Molly M. Huntsman

## Abstract

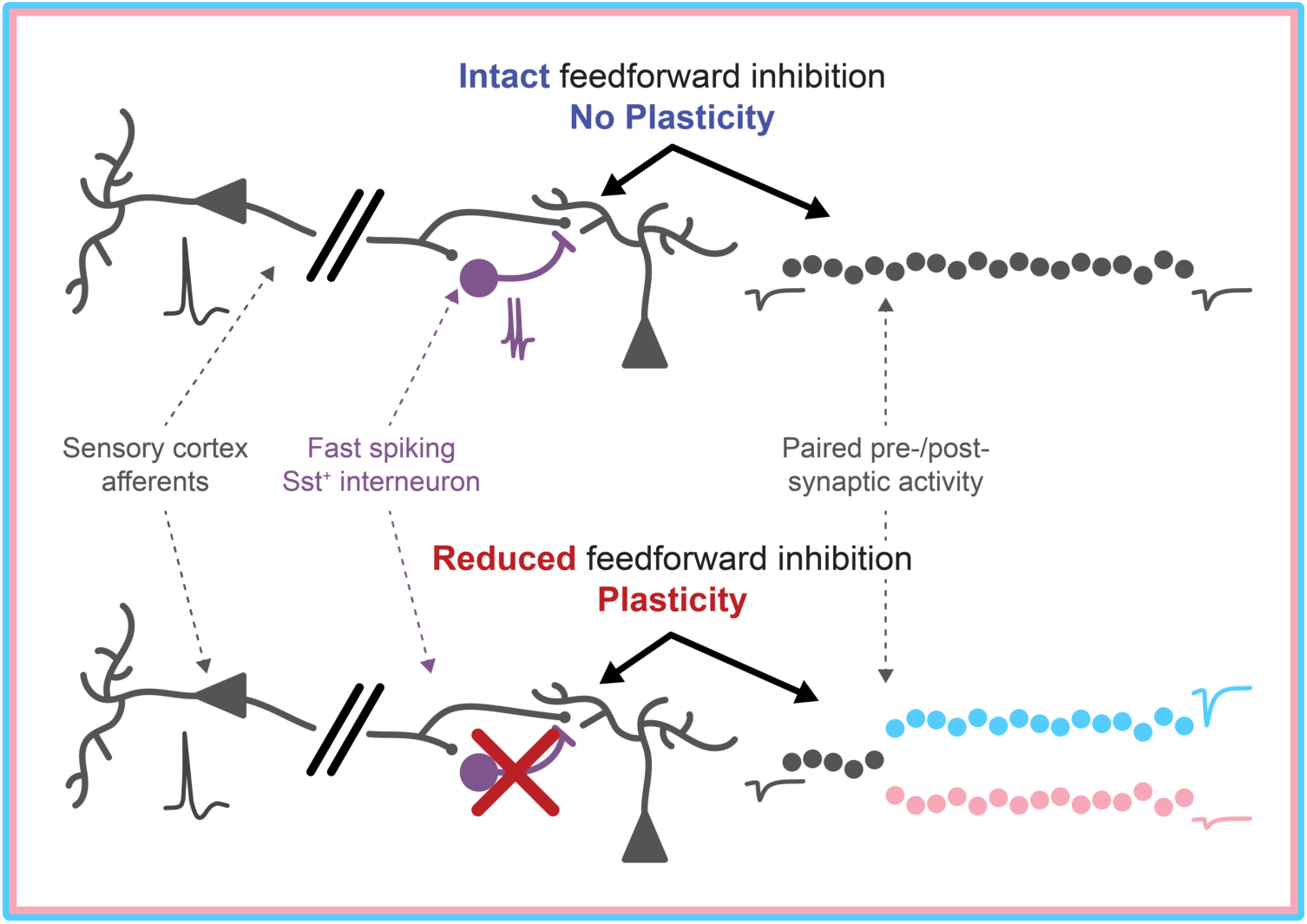

The basolateral amygdala (BLA) plays a vital role in associating sensory stimuli with salient valence information. Excitatory principal neurons (PNs) undergo plastic changes to encode this association; however, local BLA inhibitory interneurons (INs) gate PN plasticity via feedforward inhibition (FFI). Despite literature implicating parvalbumin expressing (PV^+^) INs in FFI in cortex and hippocampus, prior anatomical experiments in BLA implicate somatostatin expressing (Sst^+^) INs. The lateral entorhinal cortex (LEC), a brain region carrying olfactory information, projects to BLA where it drives FFI. In the present study, we asked whether LEC input mediates plasticity in BLA and explored the role of interneurons in this circuit. We combined patch clamp electrophysiology, chemogenetics, unsupervised cluster analysis, and predictive modeling and found that a previously unreported subpopulation of fast-spiking Sst^+^ INs mediate LEC→BLA FFI and gate plasticity. Our study raises the question whether this circuit is involved in plasticity in olfactory learning.

## INTRODUCTION

The ability of animals to learn to associate sensory stimuli with outcomes plays an important role in their survival. Specifically, animals learn to associate environmental stimuli with particular valences (rewarding or aversive outcomes). The BLA is key in the formation of these associations (Duvarci and Paré, 2014; Janak and Tye, 2015). Following a small number of presentations of a stimulus and valence information, such as a reward or threatening foot-shock, BLA excitatory PNs undergo plastic changes. For example, after a few trials BLA neurons fire selectively to novel odors that are informative about the outcome in an associative olfactory learning task (Schoenbaum et al., 1999). This plasticity is critically important as future presentations of the stimulus can then elicit robust firing in subsets of BLA PNs that output to downstream regions to guide goal-directed behavior (Beyeler et al., 2016; Janak and Tye, 2015; Quirk et al., 1995).

Accumulating evidence demonstrates that this learning induced-plasticity is input-specific (Kim and Cho, 2017; McKernan and Shinnick-Gallagher, 1997; Nabavi et al., 2014); however, the circuit mechanisms underlying this input specific plasticity remain unknown. Importantly, GABAergic inhibition regulates the ability of BLA PNs to undergo LTP with plasticity unable to occur in the presence of intact inhibition (Bissière et al., 2003; Krabbe et al., 2018). Further, local GABAergic INs expressing Sst exert strong control over BLA learning processes *in vivo*: Sst^+^ INs are normally inhibited for the duration of the sensory stimulus-valence pairing and their activation impairs learning (Wolff et al., 2014). In acute slice preparations, FFI exerts this inhibitory control over BLA plasticity (Bazelot et al., 2015; Bissière et al., 2003; Tully et al., 2007). In cerebral cortex and hippocampus, FFI is mediated by perisomatic targeting, PV^+^ INs to ensure the temporal fidelity of PN action potential (AP) output (Glickfeld and Scanziani, 2006; Pouille and Scanziani, 2001; Tremblay et al., 2016). However, in BLA, FFI plays a dual role regulating PN AP output (Lang and Paré, 1997) and gating their plasticity (Bazelot et al., 2015; Bissière et al., 2003; Tully et al., 2007). Which IN population mediates FFI to control plasticity in BLA remains unclear: perisomatic targeting PV^+^ INs are candidates as they mediate FFI in cerebral cortex and hippocampus (Glickfeld and Scanziani, 2006; Pouille and Scanziani, 2001; Tremblay et al., 2016), but anatomical studies suggest that Sst^+^ INs may play this role in the BLA (Smith et al., 2000; Unal et al., 2014). Further, the role of Sst^+^ INs in regulating BLA learning *in vivo* (Wolff et al., 2014) suggests a role in regulating BLA plasticity via FFI, and Sst^+^ INs target the dendrites and spines of BLA PNs (Muller et al., 2007; Wolff et al., 2014) placing them in prime position to regulate plasticity (Higley, 2014).

The LEC receives extensive innervation from the olfactory bulb (Igarashi et al., 2012) and piriform cortex (Johnson et al., 2000), and neurons in this area respond to odorants (Leitner et al., 2016; Xu and Wilson, 2012). Restricted firing in LEC suggests that it may play a role in modulating odor-specific, experience- and state-dependent olfactory coding (Xu and Wilson, 2012). LEC stimulation exerts an inhibitory effect on the olfactory input from the olfactory bulb to BLA and, to a lesser extent, piriform cortex (Mouly and Di Scala, 2006). Given that BLA is involved in encoding the motivational significance of olfactory cues in associative learning (Schoenbaum et al., 1999) and that the LEC→BLA synapse is plastic (Yaniv et al., 2003), the LEC→BLA circuit could be involved in plasticity during olfactory learning. In addition to its role in olfactory processing, the LEC plays an important role in multimodal sensory processing (Keene et al., 2016; Tsao et al., 2013) and the BLA is a major target of LEC efferents (McDonald, 1998). Finally, the LEC projects along the perforant pathway to engage the hippocampal trisynaptic circuit and receives input to its deep layers from CA1 (Neves et al., 2008). In turn, it is these deep layer LEC neurons that innervate the BLA (McDonald, 1998; McDonald and Mascagni, 1997). Taken together, this raises the possibility that the LEC→BLA circuit could be involved in plasticity during sensory-valence learning across sensory modalities or for more multimodal natural stimulus information under hippocampal influence. Here we address the circuit mechanisms underlying BLA dependent learning by studying the role of inhibitory interneurons in regulating the plasticity of the LEC→BLA circuit.

Due to the role of dendritic inhibition in controlling local Ca^2+^ dynamics (Chiu et al., 2013; Higley, 2014; Miles et al., 1996; Müllner et al., 2015), the role of postsynaptic Ca^2+^ dynamics in BLA plasticity (Bauer et al., 2002; Humeau et al., 2005; Humeau and Lüthi, 2007; Weisskopf et al., 1999), the importance of FFI circuits in gating BLA LTP (Bazelot et al., 2015; Bissière et al., 2003; Tully et al., 2007), and the role of BLA Sst^+^ INs in regulating learning *in vivo* (Wolff et al., 2014), we hypothesized that Sst^+^ INs provide FFI onto local PNs to gate BLA plasticity. We tested this hypothesis by pairing patch clamp recordings in BLA with unsupervised cluster analysis, predictive modeling, and chemogenetic manipulations and found that a previously unreported subpopulation of fast spiking (FS) Sst^+^ INs mediate FFI and gate plasticity in the LEC→BLA circuit.

## RESULTS

### LEC afferents preferentially drive disynaptic FFI in BLA

To examine how LEC afferents, a major cortical source of polysynaptic inhibition to BLA (Lang and Paré, 1997), engaged BLA neuronal circuitry, we prepared acute horizontal slices of mouse brain containing both BLA and LEC and recorded synaptic responses of BLA PNs (Figure 1A). A single stimulation of LEC elicited evoked EPSCs (eEPSCs) and IPSCs (eIPSCs) in PNs (Figure 1B). eEPSCs were blocked by glutamate receptor antagonists DNQX (20 μM) and D-APV (50 μM) but not by the GABA_A_ receptor antagonist gabazine (gbz; 5 μM) (Figure 1C; median [quartiles]; eEPSC_control_ = −41.17 [-34.10/-51.47] pA, eEPSC_gbz_ = −34.18 [-24.40/-39.23] pA, eEPSC_DNQX/APV_ = −4.59 [-1.49/-5.35] pA, *p* = 6.72×10^-4^, Kruskal-Wallis [KW] test, *p*_Ctrl-vs-gbz_ = 0.19, *p*_Ctrl-vs-DNQX/APV_ = 3.11×10^-4^, *p*_gbz-vs-DNQX/APV_ = 5.83×10^-4^, post-hoc Mann-Whitney U [MWU] tests with False Discovery Rate (FDR) *p*-value correction (Curran-Everett, 2000), *α*_FDR_ = 0.033, n_control_ = 8 mice, 4 cells, n_gbz_ = 7, 4, n_DNQX/APV_ = 7, 4). In contrast, eIPSCs were blocked by both gbz and DNQX/APV (Figure 1C; eIPSC_control_ = 148.82 [65.13/231.10] pA, eIPSC_gbz_ = 1.00 [-0.98/2.50] pA, eIPSC_DNQX/APV_ = 1.08 [0.56/3.15] pA, *p* = 6.45×10^-4^, KW test, *p*_Ctrl-vs-gbz_ = 3.11×10^-4^, *p*_Ctrl-vs-DNQX/APV_ = 3.11×10^-4^, *p*_gbz-vs-DNQX/APV_ = 0.71, post-hoc MWU tests, n_control_ = 8, 4; n_gbz_ = 7, 4; n_DNQX/APV_ = 7, 4), consistent with a monosynaptic glutamatergic nature of the eEPSCs and a polysynaptic GABAergic nature of the eIPSCs. Additionally, eIPSC onset was delayed relative to eEPSC onset by ∼3 ms (Figure 1D; mean ± standard error of the mean [s.e.m.]; latency_EPSC_ = 7.65 ± 0.51 ms, latency_IPSC_ = 10.72 ± 0.85 ms, *p* = 0.023, paired t-test, n = 8, 4) consistent with prior reports of polysynaptic inhibitory circuits in BLA (Arruda-Carvalho and Clem, 2014; Hübner et al., 2014; Lucas et al., 2016).

**Figure 1.**
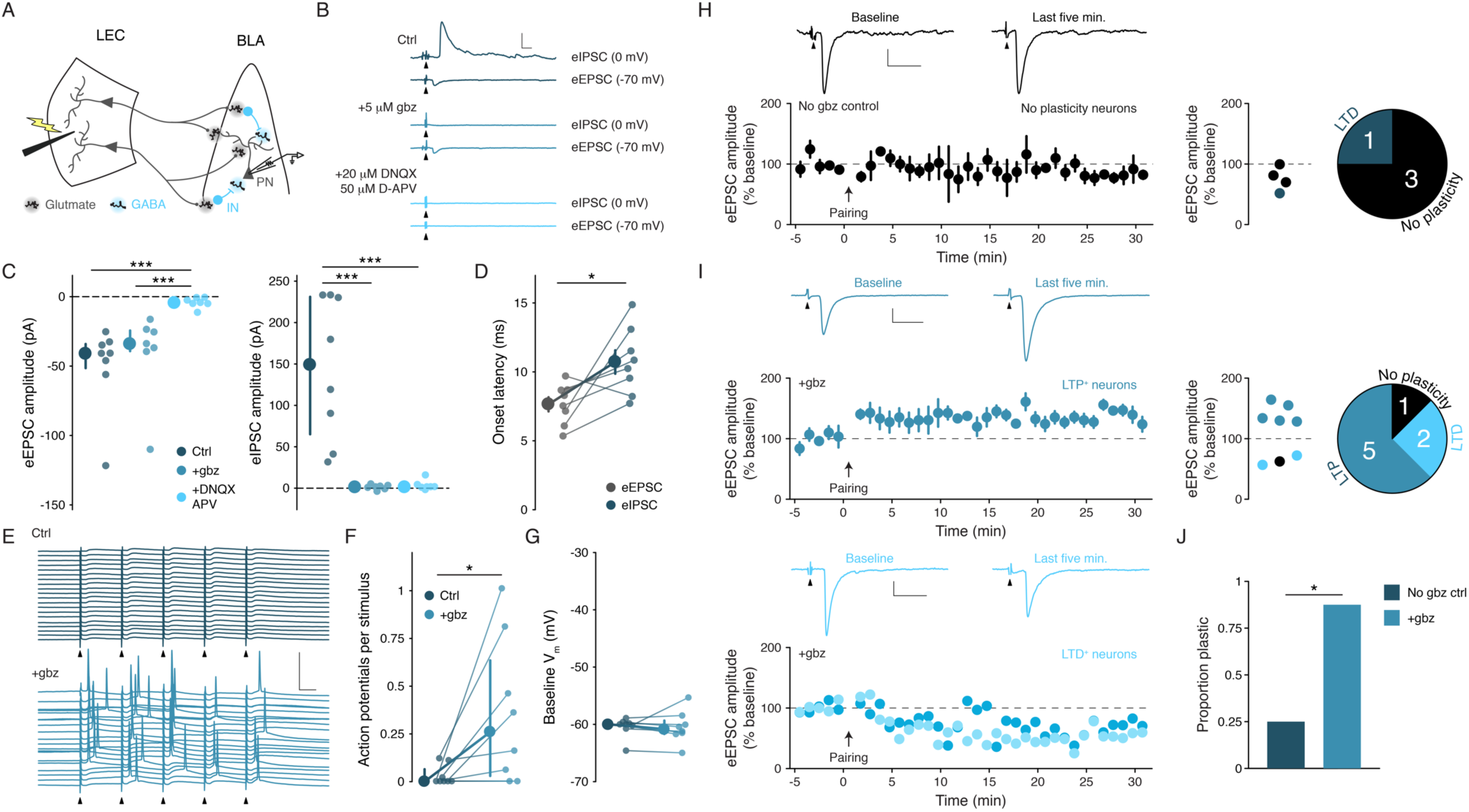
LEC afferents preferentially drive disynaptic FFI in BLA. (A) Experimental schematic. (B) Representative traces of eEPSCs and eIPSCs in a BLA PN in response to LEC stimulation (top: control; middle: gbz, 5 μM; bottom: DNQX, 20 μM, D-APV, 50 μM). Arrowheads: stimulation (artifacts truncated). Scale bars: 100 pA, 10 ms. (C) eEPSCs blocked by DNQX/APV (KW test: *p* = 6.72×10^-4^; n_control_ = 8, 4, n_gbz_ = 7, 4, n_DNQX/APV_ = 7, 4). eIPSCs blocked by gbz and DNQX/APV (KW test: *p* = 6.45×10^-4^; n_control_ = 8, 4, n_gbz_ = 7, 4, n_DNQX/APV_ = 7, 4). (D) Onset latency of eIPSC is delayed relative to the eEPSC (paired t-test: *p* = 0.023, n = 8, 4). (E) 20 voltage traces from a representative BLA PN (top: control; bottom: gbz) in response to 5× stimulation of LEC at 20 Hz. Arrowheads: stimulation. Scale bars: 40 mV, 20 ms. (F) GABA_A_ receptor blockade increases AP firing in BLA PNs in response to LEC stimulation (WSR test: *p* = 0.031, n = 8, 3). (G) Baseline V_m_ did not differ between control and gbz (WSR test: *p* = 0.38, n = 8, 3). (H) Results of pairing pre- and postsynaptic activity in the presence of intact GABAergic inhibition. Left: representative mean eEPSC trace for baseline period and last five minutes of experiment for neurons that did not undergo significant plasticity (see STAR Methods for details on significance determination; scale bars: 20 pA, 20 ms) and plot of eEPSC (% baseline) amplitude as a function of time for those neurons; data displayed as mean ± s.e.m. Arrowheads: truncated stimulus artifacts. Right: Scatter plot shows eEPSC amplitude (% baseline) in the last five minutes (n = 4, 3). Data points are colored depending on results of experiment (*p* < *α*_FDR_): significant LTD experiments or no significant plasticity. Pie chart shows results of experiments. (I) Pairing of pre- and postsynaptic activity leads to LTP and LTD in the presence of a complete GABA_A_ receptor blockade (gbz, 5 μM). Top left: representative mean eEPSC trace for baseline period and last five minutes of experiment for neurons that underwent LTP (scale bars: 50 pA, 20 ms) and plot of eEPSC (% baseline) amplitude as a function of time for those neurons; data displayed as mean ± s.e.m. Arrowheads: truncated stimulus artifacts. Bottom left: as for top left, except for LTD experiments (representative eEPSC scale bars: 20 pA, 20 ms) and plot of eEPSC amplitude as a function of time is displayed as individual neurons. Top right: Scatter plot shows eEPSC amplitude (% baseline) in the last five minutes (n = 8, 7). Data points are colored depending on results of experiment (*p* < *α*_FDR_): significant LTP experiments, significant LTD experiments, or no significant plasticity. Pie chart shows results of experiments. (J) Significantly more cells undergo long-term plastic changes during GABA_A_ receptor blockade compared to no gbz controls (X^2^ test: *p* = 0.030, n_Ctrl_ = 4, 3, n_gbz_ = 8, 7). Summary statistics in D presented as mean ± s.e.m. Summary statistics in C, F, G presented as median with interquartile range (IQR). Individual data points presented adjacent to the summary statistics. \**p* < 0.05/*α*_FDR_ (where applicable), \*\**p* < 0.01, \*\*\**p* < 0.001. See Results for post-hoc test *p* values.

Two main archetypal circuit motifs mediate polysynaptic inhibition: feedforward and feedback inhibition (Tremblay et al., 2016). In order to distinguish these motifs, we recorded from BLA PNs in response to a stimulus train of five pulses to the LEC at 20 Hz. If eIPSCs were the result of FFI alone, we would expect to see little to no AP activity in BLA PNs in response to stimulation. However, if the eIPSCs were a result of combined feedforward and feedback inhibition, we would expect to see the PNs fire in response to LEC stimulation. Recording from BLA PNs in current clamp (−60 mV), we found they rarely fired in response to stimulation (Figure 1E,F; 0.00 [0.00/0.07] APs/stimulus). These data suggest LEC stimulation preferentially drives FFI in BLA PNs.

To determine whether our experimental setup recapitulates LEC suppression of BLA PNs, as found *in vivo* (Lang and Paré, 1997), we tested the effect of gbz on the firing of BLA PNs in response to LEC stimulation (Figure 1E-G). However, GABA_A_ receptor antagonism might increase the firing of BLA PNs by depolarizing the PNs via a blockade of spontaneous or tonic inhibition. To control for this possibility, we maintained the membrane voltage (V_m_) of the PNs at −60 mV throughout these experiments (Figure 1G; V_m-control_ = −60.07 [-60.55/-59.79] mV, V_m-gbz_ = −60.91 [-61.68/-59.34] mV, *p* = 0.38, Wilcoxon signed rank [WSR] test, n = 8, 3). gbz mediated blockade of FFI led to a significant increase in PN firing in response to LEC stimulation (Figure 1E, F; APs/stimulus_control_ = 0.00 [0.00/0.07], APs/stimulus_gbz_ = 0.26 [0.03/0.64], *p* = 0.031, WSR test, n = 8, 3).

In addition to regulating PN firing (Lang and Paré, 1997), a major role of BLA FFI is to gate the induction of LTP (Bazelot et al., 2015; Bissière et al., 2003; Tully et al., 2007). To test this hypothesis, we modified a Hebbian-pairing LTP protocol used previously to study plasticity at synapses between cortical afferents and PNs of the lateral amygdala (Humeau et al., 2005). This protocol pairs EPSPs with postsynaptic APs. Each AP is followed by brief, sustained postsynaptic depolarization (25 ms) that induces backpropagating Ca^2+^ spikes (Humeau and Lüthi, 2007) and is necessary for LTP to occur (Humeau et al., 2005). For our initial experiments, we wanted to validate that this pairing protocol would induce LTP at the LEC→BLA synapse. To do this, we stimulated LEC and recorded eEPSCs in BLA PNs in the presence of 5 μM gbz to block GABA_A_ receptors. Unexpectedly, pairing presynaptic and postsynaptic activity in the presence of gbz induced a bidirectional plasticity (see STAR Methods for how direction and significance of plasticity was determined). In 7 of 8 PNs, pairing induced a long-term change in eEPSC amplitude (Figure 1I; LTP: 142.30 ± 7.33 % baseline eEPSC amplitude, n = 5, 5; LTD: 72.21 % (LTD-PN 1) and 56.71 % (LTD-PN 2), n = 2, 2). Consistent with a role in FFI in gating this plasticity, control experiments conducted in the presence of intact GABAergic inhibition showed significantly decreased plasticity at the BLA→LEC synapse (Figure 1H, J; No plasticity PNs: 83.54 ± 8.65%; n = 3, 3; Proportion plastic: control = 0.25, gbz = 0.88; *p* = 0.030, X^2^ test, n_control_ = 4, 3, n_gbz_ = 8, 7). These findings confirm that the horizontal slice preparation contained a functionally complete LEC→BLA circuit supporting its use for the study of FFI.

### Cell type specificity of the LEC→BLA circuit

We performed minimal stimulation experiments in the LEC→BLA circuit using whole cell patch clamp electrophysiology to examine putative unitary synaptic events between neurons (Gabernet et al., 2005; Kumar and Huguenard, 2001). Briefly, stimulation intensity was tuned to a threshold intensity where LEC stimulation elicits eEPSCs in BLA neurons with a ∼50% success rate and an all-or-none amplitude (Figure S1); at lower stimulation intensities eEPSCs do not occur. As these eEPSCs likely represent the response of the neuron to the activation of a single LEC neuron, this method provides a reliable measure of the unitary EPSC (uEPSC) between the stimulated and recorded neurons (Gabernet et al., 2005; Kumar and Huguenard, 2001). To identify dendritic-targeting Sst^+^ and perisomatic-targeting PV^+^ INs in acute brain slices, we generated Sst-tdTomato and PV-tdTomato mouse lines by crossing Sst-*ires*-Cre and PV-*ires*-Cre mice to *Ai9* tdTomato reporter line. Sst-tdTomato mice showed reliable labeling of Sst^+^ cells and low overlap with PV^+^ cells and PV-tdTomato mice showed reliable labeling of PV^+^ cells and low overlap with Sst^+^ cells (Figure S2). When minimally stimulating LEC we found that, whereas only a subset of BLA PNs (12/24 cells) and Sst^+^ INs (16/35 cells) responded with EPSCs, nearly every PV^+^ IN (15/16 cells) responded to the stimulation with an eEPSC (Figure 2A). DNQX and D-APV abolished eEPSCs in all cell types, confirming the glutamatergic nature of the response (Figure S1).

**Figure 2.**
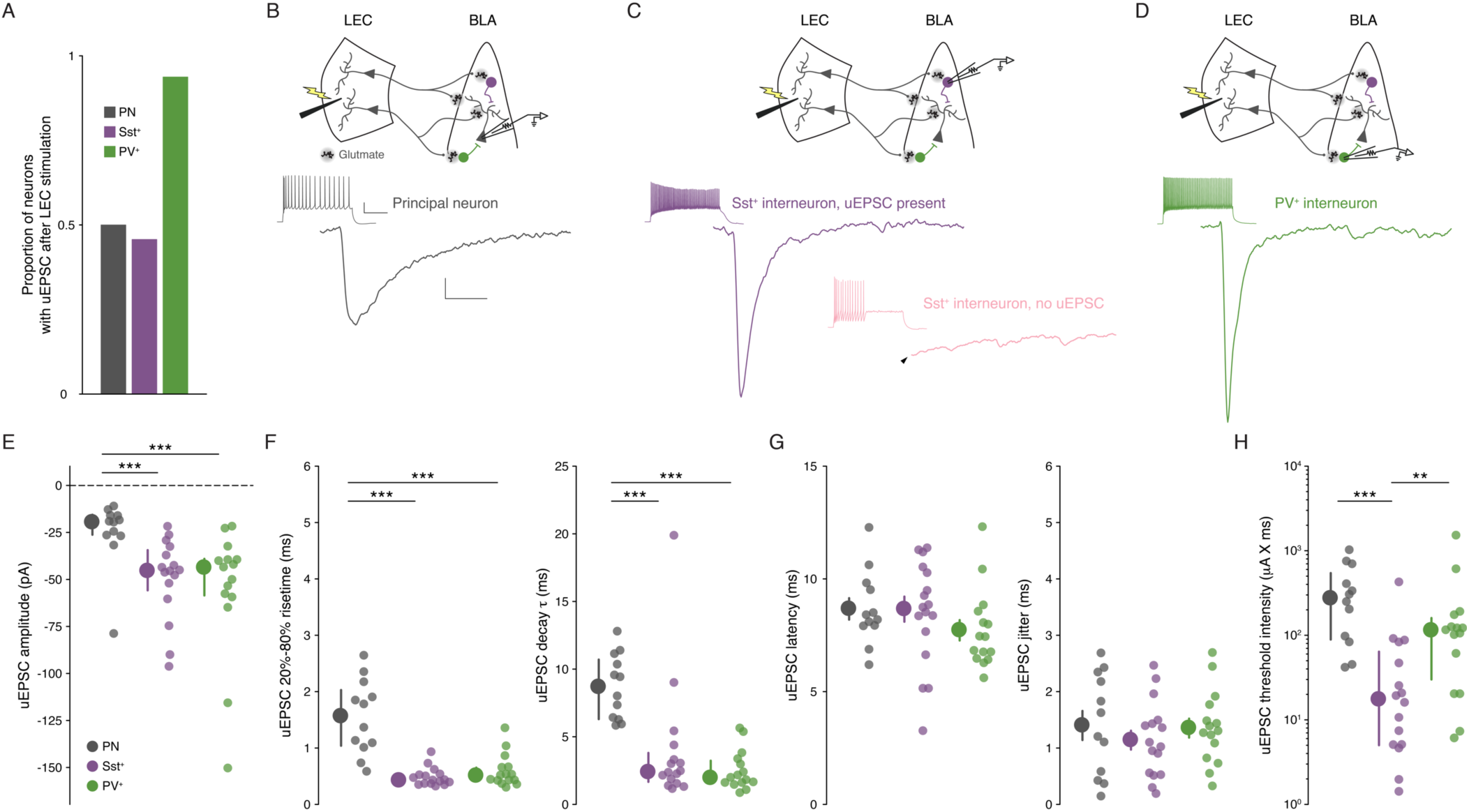
Cell type specificity of the LEC→BLA circuit. (A) Proportion of neurons with detectable uEPSCs following LEC stimulation. (B) Top: experimental schema. Bottom: mean traces of uEPSC successes from a representative PN (gray); scale bars: 5 pA, 5 ms. Inset: maximal firing to a current injection; scale bars: 20 mV, 200 ms. (C) As for B, but for a representative Sst^+^ IN with a detectable uEPSC (purple). Lower right: mean trace of lack of response in Sst^+^ IN without a detectable uEPSC (pink). Arrowhead: truncated stimulus artifact. (D) As for B, but for a representative PV^+^ IN (green). (E) uEPSC amplitude is larger in Sst^+^/PV^+^ INs compared to PNs (KW test: *p* = 5.55 × 10^-4^). (F) Left: uEPSC 20%-80% risetime is faster in Sst^+^/PV^+^ INs compared to PNs (KW test: *p* = 1.66 × 10^-5^). Right: uEPSC *τ*_Decay_ is faster in Sst^+^/PV^+^ INs compared to PNs (KW test: *p* = 2.18 × 10^-5^). (G) Left: uEPSC latency does not differ across cell types (one-way ANOVA: *p* = 0.32). Right: uEPSC jitter does not differ across cell types (one-way ANOVA: *p* = 0.60). (H) uEPSC threshold intensity is lower for Sst^+^ INs (KW test: *p* = 3.33 × 10^-4^). Summary statistics in G presented as mean ± s.e.m and in E, F, and H as median with IQR. \**p* < 0.05/*α*_FDR_ (where applicable), \*\**p* < 0.01, \*\*\**p* < 0.001. For all statistical tests: n_Sst_ = 16, 8, n_PV_ = 15, 5, n_PN_ = 12, 8. See Results for post-hoc test *p* values. See Figure S1 for example minimal stimulation experiments and data showing the glutamatergic nature of the LEC→BLA synapse. See Figure S2 for validation of the IN reporter mouse lines.

Next, we compared putative uEPSCs across cell types (Figure 2B-D). We found that the uEPSC amplitude was larger in Sst^+^ and PV^+^ INs compared to PNs (Figure 2E; Sst^+^ = −44.81 [-34.39/-55.80] pA, PV^+^ = −43.03 [-39.02/-58.51] pA, PN = −18.90 [-15.57/-26.22] pA, *p* = 5.55 × 10^-4^, KW test, *p*_Sst-vs-PV_ = 0.95, *p*_Sst-vs-PN_ = 6.45 × 10^-4^, *p*_PV-vs-PN_ = 8.30 × 10^-4^, post-hoc MWU tests, *α*FDR = 0.033, n_Sst_ = 16, 8, n_PV_ = 15, 5, n_PN_ = 12, 8). Additionally, we found the uEPSC kinetics were faster in the INs compared to PNs (Figure 2F; 20%-80% risetime: Sst^+^ = 0.42 [0.36/0.52] ms, PV^+^ = 0.51 [0.41/0.65] ms, PN = 1.56 [1.04/2.03] ms, *p* = 1.66 × 10^-5^, KW test, *p*_Sst-vs-PV_ = 0.29, *p*_Sst-vs-PN_ = 2.16 × 10^-5^, *p*_PV-vs-PN_ = 1.56 × 10^-4^, post-hoc MWU tests, *α*FDR = 0.033; *τ*_Decay_: Sst^+^ = 2.38 [1.67/3.80] ms, PV^+^ = 1.95 [1.57/3.23] ms, PN = 8.68 [6.30/10.70] ms, *p* = 2.18 × 10^-5^, KW test, *p*_Sst-vs-PV_ = 0.40, *p*_Sst-vs-PN_ = 3.21 × 10^-4^, *p*_PV-vs-PN_ = 1.26 × 10^-5^, post-hoc MWU tests, *α*FDR = 0.033, n_Sst_ = 16, 8, n_PV_ = 15, 5, n_PN_ = 12, 8). Finally, we found no differences between cell types in uEPSC latency or jitter (Figure 2G; latency: Sst^+^ = 8.66 ± 0.56 ms, PV^+^ = 7.72 ± 0.45 ms, PN = 8.67 ± 0.48 ms, *p* = 0.32, one-way ANOVA; jitter: Sst^+^ = 1.14 ± 0.17 ms, PV^+^ = 1.35 ± 0.17 ms, PN = 1.40 ± 0.26 ms, *p* = 0.60, one-way ANOVA, n_Sst_ = 16, 8, n_PV_ = 15, 5, n_PN_ = 12, 8). Together with our data demonstrating that BLA PNs rarely fire in response to LEC stimulation (Figure 1E, F), the statistically indistinguishable low jitters are consistent with a monosynaptic connection between LEC projection neurons and BLA Sst^+^ INs, PV^+^ INs, and PNs (Doyle and Andresen, 2001).

Since minimal stimulation likely reports the response of a BLA neuron to a single upstream LEC neuron (Gabernet et al., 2005; Kumar and Huguenard, 2001), we can use threshold stimulation intensity as an indirect measure of convergence of LEC projection neurons onto different BLA cell types. If threshold stimulation is lower for one BLA neuronal population compared to the others, it would follow that convergence of LEC inputs onto that cell type is likely greater relative to the other populations as it requires the activation of fewer LEC neurons. We found that threshold intensity was lower for the Sst^+^ INs compared to other cell types (Figure 2H; Sst^+^ = 17.40 [5.01/64.00] μA×ms, PV^+^ = 114.00 [30.00/160.00] μA×ms, PN = 273.00 [89.00/544.00] μA×ms, *p* = 3.33 × 10^-4^, KW test, *p*_Sst-vs-PV_ = 0.0047, *p*_Sst-vs-PN_ = 3.83 × 10^-4^, *p*_PV-vs-PN_ = 0.092, post-hoc MWU tests, *α*FDR = 0.033, n_Sst_ = 16, 8, n_PV_ = 15, 5, n_PN_ = 12, 8).

Taken together, these data demonstrate that LEC stimulation leads to large and fast unitary currents in BLA Sst^+^ and PV^+^ INs and small and slow unitary currents in BLA PNs. Finally, although Sst^+^ and PV^+^ INs display equivalent uEPSCs, the finding that Sst^+^ INs have a lower threshold stimulation intensity compared to PV^+^ INs and PNs suggest that LEC afferents may have a greater functional convergence onto BLA Sst^+^ compared to PV^+^ INs.

### A fast spiking phenotype distinguishes BLA Sst^+^ INs targeted by LEC afferents

Though little is known about BLA Sst^+^ INs, they appear to have diverse electrophysiological properties *ex vivo* (Krabbe et al., 2018; Sosulina et al., 2010) and responses to stimuli *in vivo* (Krabbe et al., 2018; Wolff et al., 2014), suggesting Sst may be expressed by a broad range of GABAergic IN subtypes, similar to cerebral cortex (Tremblay et al., 2016). Only a subset of Sst^+^ INs responded to cortical stimulation raising the question whether these cells represented a distinct cell type. To address this hypothesis, we ran an unsupervised cluster analysis using Ward’s method (Ward, 1963) based on 15 membrane properties from 105 Sst^+^ INs. Applying Thorndike’s procedure (Thorndike, 1953) suggested two distinct clusters (Groups I and II). A large majority of Sst^+^ INs that responded to cortical stimulation clustered into Group I (14/16 cells) whereas a large majority of non-responsive Sst^+^ INs clustered into Group II (17/19 cells) (Figure 3A). To better understand what membrane properties best distinguished Group I and II Sst^+^ INs, we used decision tree analysis (Breiman et al., 1984; Therneau and Atkinson, n.d.) which returned maximum firing rate, hyperpolarization induced sag, and AHP latency as the most salient parameters for cluster separation (Figure S3). To reduce potential model overfitting with the decision tree analysis, we used an additional predictive modeling technique, the random forest method (Liaw and Wiener, 2002). Like decision tree analysis, the random forest method returned maximum firing rate and hyperpolarization induced sag in addition to AP halfwidth as the most salient parameters for defining BLA Sst^+^ INs (Figure 3B, C).

**Figure 3.**
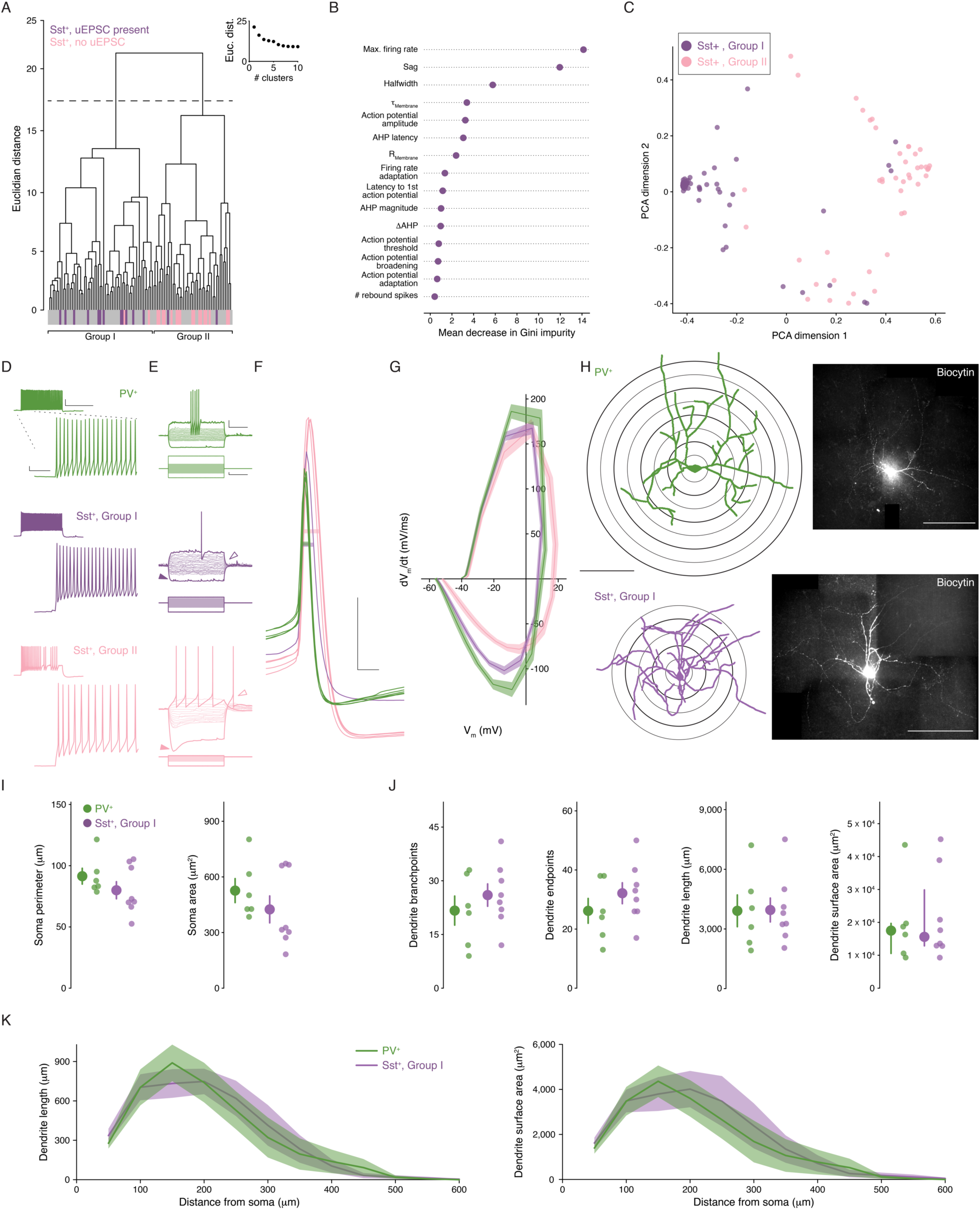
A fast spiking phenotype distinguishes BLA Sst^+^ INs targeted by LEC afferents. (A) Unsupervised clustering analysis revealed two clusters of Sst^+^ INs. Majority of Sst^+^ INs with a uEPSC present following LEC stimulation cluster in Group I (14/16, Sst^+^ INs with uEPSC; 60/105, all Sst^+^ INs). Majority of Sst^+^ INs with no response to LEC stimulation cluster in Group II (17/19, Sst^+^ INs without uEPSC; 45/105, all Sst^+^ INs). (B) Results of Random Forest model. Plot shows mean decrease in Gini impurity for each of the 15 parameters included in the model (n_Sst_ = 105, 23). Gini impurity measures how often a random Sst^+^ IN would be clustered incorrectly if labeled randomly according to the distribution of labels in the set. (C) First two dimensions of the principal components analysis (PCA) of the random forest proximity matrix (n_Sst_ = 105, 23). (D) Maximum AP firing from representative PV^+^ (top, green), Group I (middle, purple) and II (bottom, pink) Sst^+^ INs. Scale bars, main traces: 5 mV, 50ms; inset: 20 mV, 400 ms. (E) Voltage responses of the same representative PV^+^ (top, green), Group I (middle, purple) and II (bottom, pink) Sst^+^ INs. Darker traces show responses to −200 pA and rheobase current injections. Lighter traces show responses to intermediate current injections were used to determine membrane resistance (−100 pA to sweep immediately before rheobase, *Δ*10 pA each sweep, +100 pA maximum current injection). Note lack of sag, rebound AP in Group I Sst^+^ IN (sag: closed arrowheads; rebound AP: open arrowheads). Scale bars: 10 mV, 200 ms; 50 pA, 200 ms. (F) APs of the same representative PV^+^ (green), Group I (purple), and II (pink) Sst^+^ INs at rheobase. Halfwidth: bar through width of AP. Scale bars: 20 mV, 1 ms. (G) Phase plots of PV^+^, Group I, and Group II Sst^+^ INs shows rate of voltage change for Group I and II APs at rheobase. Data presented as mean ± s.e.m. n_PV_ = 52, 19, n_SstI_ = 60, 23, n_SstII_ = 45, 18. (H) Left: reconstructed soma and dendrites of a representative PV^+^ and Group I Sst^+^ INs (scale bar: 200 μm). Sholl rings shown beneath reconstruction. Right: confocal images of the biocytin filled PV^+^ and Group I Sst^+^ INs (scale bar: 200 μm). (I) No significant differences between PV^+^ and Group I Sst^+^ INs for soma perimeter (left; unpaired t-test: *p* = 0.26) or area (right; MWU test: *p* = 0.28). (J) No significant differences between PV^+^ and Group I Sst^+^ INs for (left to right) dendrite branchpoints (unpaired t-test: *p* = 0.40), endpoints (unpaired t-test: *p* = 0.30), length (unpaired t-test: *p* = 0.97), or surface area (MWU test: *p* = 0.95). (K) Results of Sholl analysis showing dendrite length (left) and surface area (right) as a function of distance from the soma in PV^+^ and Group I Sst^+^ INs. Data presented as mean ± s.e.m. n_PV_ = 6, 4, n_SstI_ = 8, 6. Summary statistics in I (perimeter) and J (branchpoints, endpoints, and length) presented as mean ± s.e.m. Summary statistics in I (area) and J (area) presented as median and IQR. For all statistical tests in (I) and (J): n_PV_ = 6, 4, n_SstI_ = 8, 6. See Figure S3 for decision tree data and S4 for data on Group II Sst^+^ IN morphology. See Table S1 for data on all membrane properties studied for Group I and Group II Sst^+^ INs and PV^+^ INs.

Having observed two subpopulations of Sst^+^ INs, we compared membrane properties across Group I and II Sst^+^ INs and PV^+^ INs (Figure 3D-G, Table S1). These data show that Group I Sst^+^ and PV^+^ INs differed from Group II Sst^+^ INs in a subset of membrane properties such as maximum firing rate and hyperpolarization induced sag. Further, all three IN subtypes differed in AP halfwidth with PV^+^ INs firing the fastest APs and Group I Sst^+^ INs firing faster APs than Group II Sst^+^ INs (see Table S1 for detailed statistics on membrane properties). The membrane properties of Group I Sst^+^ and PV^+^ INs are consistent with a FS phenotype typically seen in cortical, hippocampal, and BLA PV^+^ INs that is characterized by the ability to fire high frequency trains of brief APs (Tremblay et al., 2016; Woodruff and Sah, 2007a; however, see [Large et al., 2016; Ma et al., 2006; Nigro et al., 2018] for examples of FS Sst^+^ INs in cerebral cortex).

In order to compare the morphology of the IN populations, we included biocytin in the patch pipette in a subset of recordings to allow for *post hoc* visualization of the different IN subtypes. Using these biocytin-filled neurons, we created reconstructions of the somatic and dendritic morphology of the INs (Figure 3H; see Figure S4 for Group II Sst^+^ IN data which were not included in formal analyses due to low number of recovered morphologies [n_SstII_ = 3, 2]). When we compared the somatic and dendritic morphology of PV^+^ and Group I Sst^+^ INs, we found no significant differences in any measurement (Figure 3I-K; soma perimeter: PV^+^ = 91.42 ± 6.43 μm, Group I Sst^+^ = 79.96 ± 6.95 μm, *p* = 0.26, unpaired t-test; soma area: PV^+^ = 463.87 [426.50/615.37] μm^2^, Group I Sst^+^ = 320.68 [287.76/663.36] μm^2^, *p* = 0.28, MWU test; dendrite branchpoints: PV^+^ = 21.67 ± 4.05, Group I Sst^+^ = 26.00 ± 3.11, *p* = 0.40, unpaired t-test; dendrite endpoints: PV^+^ = 26.17 ± 4.18, Group I Sst^+^ = 32.13 ± 3.53, *p* = 0.30, unpaired t-test; dendrite length: PV^+^ =3,911.09 ± 797.02 μm, Group I Sst^+^ = 3,952.67 ± 598.47 μm, *p* = 0.97, unpaired t-test; dendrite surface area: PV^+^ = 1.75 × 10^%^ [1.06 × 10^%^/1.97 × 10^%^] μm^2^, Group I Sst^+^ = 1.56 × 10^%^ [1.29 × 10^%^/2.99 × 10^%^] μm^2^, *p* = 0.95, MWU test; n_PV_ = 6, 4, n_SstI_ = 8, 6).

Taken together, the membrane properties of BLA Sst^+^ and PV^+^ INs reveal two distinct subpopulations of Sst^+^ INs that are readily distinguished at the biophysical level by their FS phenotype and at the functional circuit level by synaptic responses to cortical stimulation. However, the IN subtypes do not appear to have any readily observable differences in their somatic and dendritic morphology. For clarity, we refer to the Group I and II Sst^+^ INs as FS and non-fast spiking (nFS) Sst^+^ INs, respectively.

### Probing the LEC→BLA circuitry suggests distinct functional feedforward/feedback roles for IN subtypes

BLA Sst^+^ INs have lower threshold stimulation intensity compared to PV^+^ INs and PNs (Figure 2H). Since these data suggest a higher rate of convergence of LEC afferents onto Sst^+^ compared to PV^+^ INs, we wanted to test the hypothesis that LEC input to BLA may preferentially recruit Sst^+^ over PV^+^ INs. To do this, we recorded from BLA Sst^+^ and PV^+^ INs in current clamp at rest and stimulated LEC with five pulses at 20 Hz. The stimulation intensity was set to the empirically derived threshold stimulation for BLA PNs (defined as the median PN threshold stimulation; 273 μA×ms; Figure 2H) allowing us to determine how BLA INs respond when LEC activity is sufficient to ensure that PNs receive input. We found that stimulation led to robust spiking in BLA Sst^+^ INs whereas PV^+^ INs rarely fired (Figure 4A, B; APs/stimulus: Sst^+^ = 0.93 [0.78/1.01], PV^+^ = 0.00 [0.00/0.003], *p* = 0.016, MWU test, n_Sst_ = 8, 3, n_PV_ = 9, 3).

**Figure 4.**
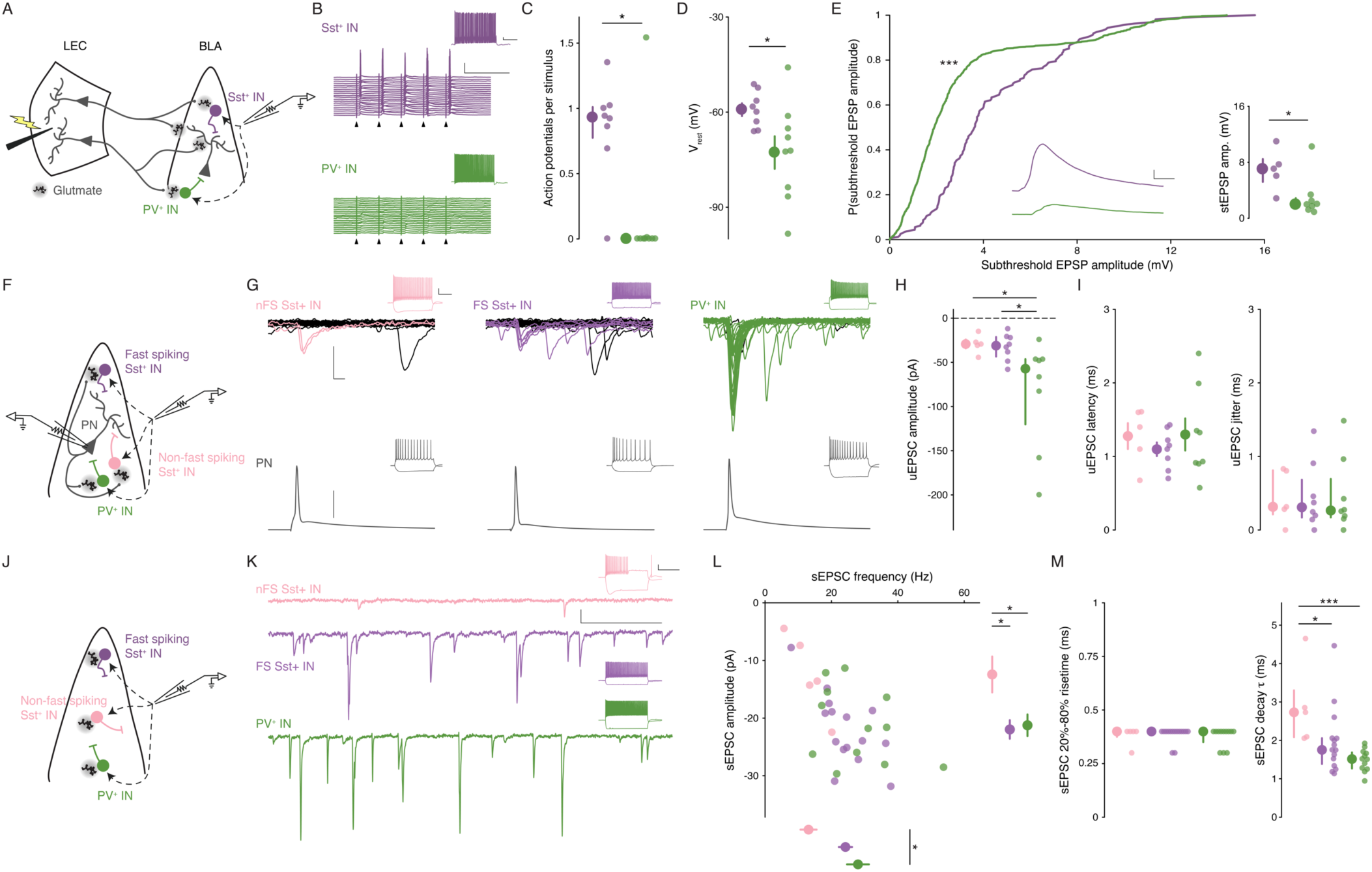
Distinct microcircuit functional roles for IN subtypes. (A) Experimental schematic for B-E. (B) Representative Sst^+^ (top, purple) and PV^+^ (bottom, green) responses to LEC stimulation at 20 Hz (arrowhead: stimulation artifacts; scale bars: 25 mV, 100 ms). Insets show maximal firing frequency for the representative IN in response to a square current pulse (scale bars: 5 mV, 200 ms). (C) Sst^+^ INs fire more APs per stimulus compared to PV^+^ INs (MWU test: *p* = 0.016; n_Sst_ = 8, 3; n_PV_ = 9, 3). (D) Sst^+^ INs that respond to LEC stimulation have a more depolarized V_rest_ compared to PV^+^ INs (unpaired t-test: *p* = 0.034; n_Sst_ = 8, 3; n_PV_ = 9, 3). (E) Cumulative probability distribution of all stEPSPs. Inset, middle: mean stEPSP from representative neurons in (A; Sst^+^: purple, PV^+^: green; scale bars: 1 mV, 5 ms; KS test: *p* = 2.19 × 10^-21^, n_Sst_ = 161 events, n_PV_ = 779 events; MWU test: *p* = 0.030, n_Sst_ = 5, 3, n_PV_ = 8, 3). (F) Experimental schematic for G-I. (G) Representative paired recording experiments between BLA and INs (left to right: nFS Sst^+^, FS Sst^+^, PV^+^). Bottom: AP in BLA PN (scale bar: 40 mV). Top: overlaid current responses of BLA INs to PN AP (50 pA, 2ms); successful trials shown in color. Insets show maximal firing frequency and response to a −200 pA current injection in the representative neurons (scale bars: 20 mV, 200 ms). (H) uEPSC amplitude is larger in PV^+^ compared to FS and nFS Sst^+^ INs (KW test: *p* = 0.025, n_nFS-Sst_ = 5, 5, n_FS-Sst_ = 8, 6, n_PV_ = 8, 8) (I) No significant differences in uEPSC latency or jitter across IN subtypes (latency, one-way ANOVA: *p* = 0.65, n_nFS-Sst_ = 5, 5, n_FS-Sst_ = 8, 6, n_PV_ = 8, 8; jitter, KW test: *p* = 0.97, n_nFS-Sst_ = 5, 5, n_FS-Sst_ = 8, 6, n_PV_ = 8, 8). (J) Experimental schematic for K-M. (K) Representative sEPSC traces from BLA INs (top to bottom: nFS Sst^+^, FS Sst^+^, PV; scale bars: 20 pA, 100 ms). Insets show maximal firing frequency and response to a −200 pA current injection in the representative neurons (scale bars: 20 mV, 300 ms). (L) Scatter plot of sEPSC frequency and amplitude. Right: nFS Sst^+^ INs have smaller amplitude sEPSCs compared to FS Sst^+^ and PV^+^ INs (one-way ANOVA: *p* = 0.022). Bottom: PV^+^ INs have more frequent sEPSCs compared to nFS Sst^+^ INs (one-way ANOVA: *p* = 0.022). (M) Left: no significant differences in sEPSC 20%-80% risetime (KW test: *p* = 0.75). Right: nFS Sst^+^ IN sEPSCs have slower decay kinetics compared to FS Sst^+^ and PV^+^ IN sEPSCs (KW test: *p* = 0.0033). Summary statistics in D, I (left), and L presented in color as mean ± s.e.m. and in C, E (inset), H, I (right), and M in color as median with IQR. \**p* < 0.05, \*\*\**p* < 0.001. See Results for post-hoc test *p* values.

To probe the underlying mechanisms of the preferential spiking of Sst^+^ INs, we used two additional measures: resting V_m_ (V_rest_) and subthreshold EPSP (stEPSP) amplitude. We found that V_rest_ was more depolarized in Sst^+^ compared to PV^+^ INs (Figure 4C; V_rest_: Sst^+^ = −59.24 ± 2.01 mV, PV^+^ = −72.76 ± 5.14 mV, *p* = 0.034, unpaired t-test, n_Sst_ = 8, 3, n_PV_ = 9, 3). Because individual INs displayed variability in the number of APs/stimulus, each IN would have a different number of stEPSPs. Therefore, we compared both the distribution of all stEPSP amplitudes and the mean cellular stEPSP amplitudes across Sst^+^ and PV^+^ INs. Regardless of the analysis method, we found that evoked stEPSPs were larger in Sst^+^ compared to PV^+^ INs (Figure 4D; all stEPSP events: *p* = 2.19 × 10^-21^, Kolmogorov-Smirnov [KS] test, n_Sst_ = 161 events, n_PV_ = 779 events; cellular stEPSP: Sst^+^ = 7.01 [5.19/8.46] mV, PV^+^ = 1.95 [1.41/2.87] mV, *p* = 0.030, MWU test, n_Sst_ = 5, 3, n_PV_ = 8, 3). Thus, the data show that two distinct mechanisms underlie the recruitment of BLA Sst^+^ INs by LEC afferents: Sst^+^ INs are more depolarized at rest compared to PV^+^ INs, and, despite the concomitant decrease in the driving force through glutamate receptors, the magnitude of evoked stEPSPs was greater in Sst^+^ compared to PV^+^ INs. Finally, when we classified the Sst^+^ INs from this experiment (Figure 3A), we found that all but one (7/8 cells) were FS Sst^+^ INs, consistent with our finding that LEC afferents appear to selectively target FS cells among the Sst^+^ INs.

Taken together with the data in Figures 2 and 3, these data show that LEC stimulation leads to synaptic responses in BLA FS Sst^+^ INs, PV^+^ INs, and PNs but not in nFS Sst^+^ INs. Although FS Sst^+^ and PV INs have equivalent unitary events following LEC stimulation, the responses of these cell types to LEC input diverge in important ways. Specifically, FS Sst^+^ INs have a lower threshold stimulation intensity compared to PV^+^ INs, and FS Sst^+^ INs have greater stEPSP amplitudes and are more likely to fire in response to PN threshold stimulation of LEC compared to PV^+^ INs. These findings suggest that LEC afferents to BLA have a greater functional convergence onto FS Sst^+^ INs compared to PV^+^ INs and raise the question whether they are involved in BLA FFI.

In neocortical circuits, PV^+^ INs mediate FFI whereas Sst^+^ typically mediate feedback inhibition (Tremblay et al., 2016); nevertheless, our data point to a role for Sst^+^ INs in BLA FFI, consistent with prior hypotheses that BLA Sst^+^ INs mediate FFI and PV^+^ INs mediate feedback inhibition (Duvarci and Paré, 2014). Thus, we wanted to test for responses of Sst^+^ and PV^+^ INs to local BLA PNs to probe for potential roles in feedback inhibition (Figure 4F). To do this, we recorded PN-IN pairs in BLA, drove AP firing in PNs and compared uEPSC responses to the AP across IN subtypes (Figure 4G). We found responses in all IN subtypes (8/10 FS Sst^+^, 5/5 nFS Sst^+^, 8/12 PV^+^ INs with uEPSCs in response to PN APs) and found that BLA PV^+^ INs responded to PNs with larger amplitude uEPSCs compared to either FS or nFS Sst^+^ INs (Figure 4H; nFS Sst^+^ = −29.30 [-24.15/-34.16] pA, FS Sst^+^ = −31.13 [-21.34/-43.43] pA, PV^+^ = −57.16 [-46.38/-120.23] pA, *p* = 0.025, KW test, *p*_nFS-Sst-vs-FS-Sst_ = 0.72, *p*_nFS-Sst-vs-PV_ = 0.019, *p*_FS-Sst-vs-PV_ = 0.028, post-hoc MWU tests, *α*FDR = 0.033, n_nFS-Sst_ = 5, 5, n_FS-Sst_ = 8, 6, n_PV_ = 8, 8). Further, we found no difference in the latency or jitter of the uEPSCs across cell types (Figure 4I; latency: nFS Sst^+^ = 1.28 ± 0.18 ms, FS Sst^+^ = 1.10 ± 0.092 ms, PV^+^ = 1.29 ± 0.22 ms, *p* = 0.65, one-way ANOVA; jitter: nFS Sst^+^ = 0.31 [0.17/0.69] ms, FS Sst^+^ = 0.31 [0.17/0.68] ms, PV^+^ = 0.26 [0.21/0.81] ms, *p* = 0.97, KW test, n_nFS-Sst_ = 5, 5, n_FS-Sst_ = 8, 6, n_PV_ = 8, 8). These data are consistent with a larger role for PV^+^ INs in BLA feedback inhibition relative to Sst^+^ INs.

Finally, we wanted to look at the level of spontaneous excitation onto the different IN subtypes as different levels of spontaneous glutamatergic activity could lead to different levels of basal excitability for the INs in the LEC→BLA circuit (Figure 4J, K). We recorded spontaneous EPSCs (sEPSCs) across BLA IN subtypes. We found that PV^+^ INs had larger amplitude sEPSCs than either Sst^+^ IN subtype and that PV^+^ and FS Sst^+^ INs had more frequent sEPSCs than nFS Sst^+^ Ins (Figure 4L; amplitude: nFS Sst^+^ = 12.43 ± 3.12 pA, FS Sst^+^ = 21.97 ± 1.61 pA, PV^+^ = 21.24 ± 1.88 pA, *p* = 0.022, one-way ANOVA, *p*_nFS-Sst-vs-FS-Sst_ = 0.021, *p*_nFS-Sst-vs-PV_ = 0.041, *p*_FS-Sst-vs-PV_ = 0.95; frequency: nFS Sst^+^ = 13.10 ± 2.43 Hz, FS Sst^+^ = 24.21 ± 2.00 Hz, PV^+^ = 27.99 ± 3.29 Hz, *p* = 0.022, one-way ANOVA, *p*_nFS-Sst-vs-FS-Sst_ = 0.061, *p*_nFS-Sst-vs-PV_ = 0.012, *p*_FS-Sst-vs-PV_ = 0.54, n_nFS-Sst_ = 5, 5, n_FS-Sst_ = 15, 8, n_PV_ = 12, 5). Additionally, sEPSCs in nFS Sst^+^ INs had slower decay kinetics compared to the other BLA IN subtypes (Figure 4M; 20%-80% risetime: nFS Sst^+^ = 0.40 [0.38/0.40] ms, FS Sst^+^ = 0.40 [0.40/0.40] ms, PV^+^ = 0.40 [0.35/0.40] ms, *p* = 0.75, KW test; *τ*_Decay_: nFS Sst^+^ = 2.73 [2.08/3.30] ms, FS Sst^+^ = 1.75 [1.39/2.06] ms, PV^+^ = 1.51 [1.27/1.69] ms, *p* = 0.0033, KW test, *p*_nFS-Sst-vs-FS-Sst_ = 0.023, *p*_nFS-Sst-vs-PV_ = 3.23 × 10^-4^, *p*_FS-Sst-vs-PV_ = 0.083, post-hoc MWU tests, *α*FDR = 0.033, n_nFS-Sst_ = 5, 5, n_FS-Sst_ = 15, 8, n_PV_ = 12, 5). These data suggest that although FS Sst^+^ and PV^+^ INs may have divergent functional roles with regards to feedforward and feedback inhibition, they both receive greater levels of spontaneous excitatory input compared to nFS Sst^+^ INs.

### Sst^+^ INs mediate LEC→BLA FFI

To test whether Sst^+^ or PV^+^ INs provide cortically evoked FFI onto BLA PNs, we generated Sst-hM4Di and PV-hM4Di mice by crossing the Sst-*ires*-Cre or PV-*ires*-Cre mice to the ROSA-hM4Di-mCitrine mice. These mice selectively express hM4Di in Sst^+^ (Sst-hM4Di) or PV^+^ (PV-hM4Di) INs. hM4Di is an inhibitory chemogenetic receptor that, when bound to its ligand clozapine-*N*-oxide (CNO), activates the G_i/o_ signaling cascade to hyperpolarize and block neurotransmitter release from neurons (Armbruster et al., 2007; Stachniak et al., 2014). To validate the efficacy of the Sst-hM4Di and PV-hM4Di mouse lines, we patched onto BLA mCitrine^+^ neurons and recorded V_m_ in current clamp (I_hold_ = 0 pA). Application of 10 μM CNO reduced V_m_ of mCitrine^+^ neurons in both lines (Figure 5A; *Δ*V_m-Sst_ = −7.18 ± 2.21 mV, *Δ*V_m-PV_ = −9.43 ± 3.16 mV, *p*_Sst_ = 0.017, *p*_PV_ = 0.031, one-sample t-test, n_Sst_ = 7, 4, n_PV_ = 6, 3). Thus, we used the Sst-hM4Di and PV-hM4Di mice to selectively hyperpolarize Sst^+^ and PV^+^ INs.

**Figure 5.**
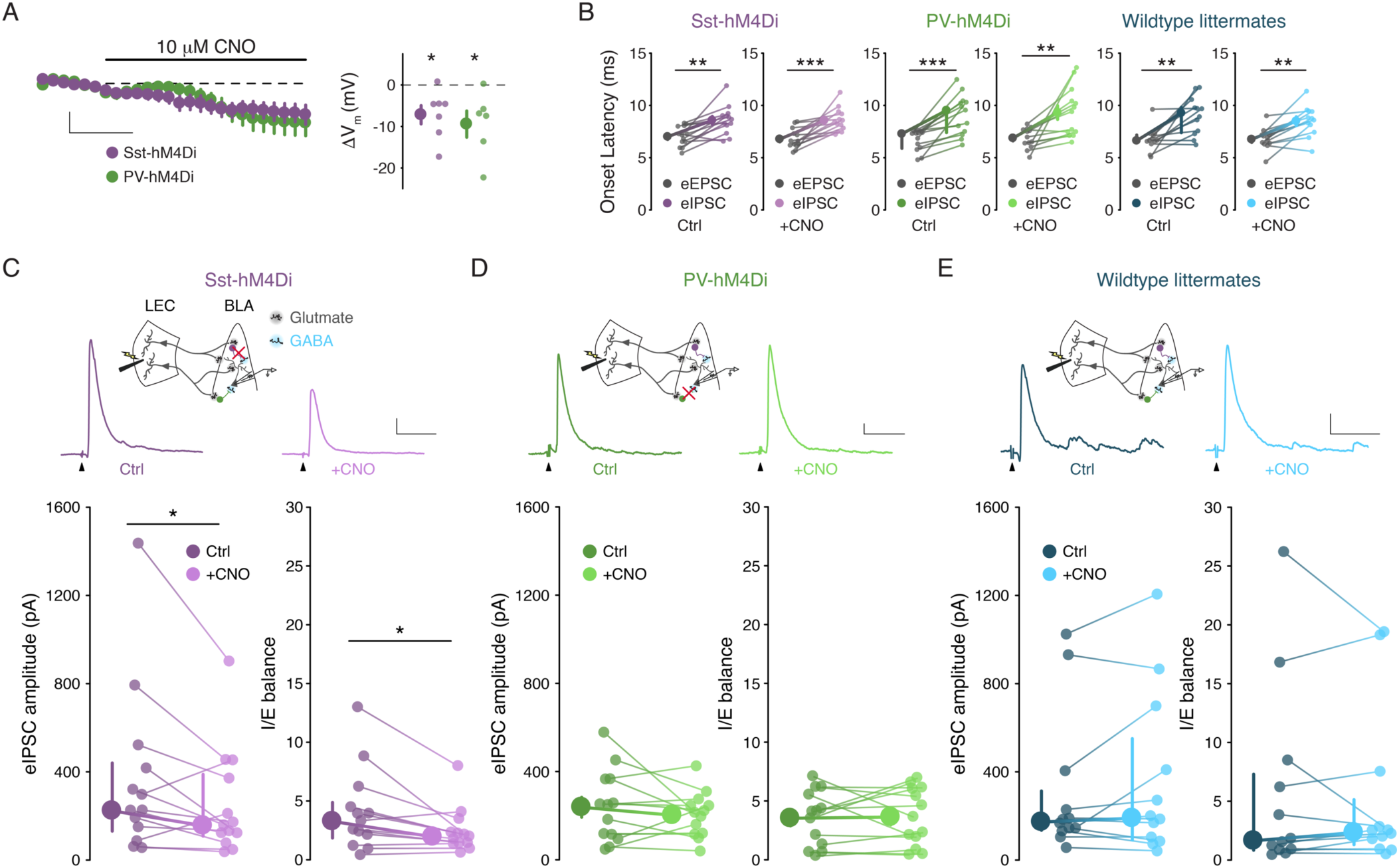
Sst^+^ INs mediate LEC→BLA FFI. (A) Left: Effect of CNO on V_m_. Scale bars: 5 mV, 3 min. Right: CNO hyperpolarizes mCitrine^+^ cells in both mouse lines (*p*_Sst_ = 0.017, *p*_PV_ = 0.031, one-sample t-test, n_Sst_ = 7, 4, n_PV_ = 6, 3). (B) eIPSC is delayed relative to eEPSC (paired t-tests: *p*_Sst-Ctrl_ = 0.0018, *p*_Sst-CNO_ = 8.80 × 10^-4^, *p*_PV-CNO_ = 0.0042, *p*_WT-CNO_ = 0.0012; WSR tests: *p*_PV-Ctrl_ = 2.44 × 10^-4^, *p*_WT-Ctrl_ = 0.0024; n_Sst_ = 13, 6, n_PV_ = 13, 7, n_WT_ = 12, 6). (C) CNO mediated Sst^+^ IN inactivation reduces FFI (Sst-hM4Di mouse line; WSR tests: *p*_IPSC_ = 0.040, *p*_IE_ = 0.013, n = 13, 6). Top: Experimental schema, eIPSC traces with and without CNO. Arrowheads: stimulation artifacts (truncated). Scale bars: 100 pA, 50ms. (D) PV^+^ IN inactivation has no effect on FFI (paired t-tests: *p*_IPSC_ = 0.35, *p*_IE_ = 0.85, n = 13, 7). Display as for (C), but in PV-hM4Di mouse line. Scale bars: 25 pA, 50ms. (E) CNO has no effect on FFI in wildtype littermates (WSR tests: *p*_IPSC_ = 0.73, *p*_IE_ = 0.68, n = 12, 6). Display as for (C), but in hM4Di^-/-^ littermates of Sst- and PV-hM4Di mice. Scale bars: 50 pA, 50 ms. Summary statistics in A, B (Sst-hM4Di both conditions, PV-hM4Di CNO condition, WT littermates CNO condition), and D in color as mean ± s.e.m. Summary statistics in B (PV-hM4Di and WT littermates control conditions), C, and E in color as median with IQR. \**p* < 0.05, \*\**p* < 0.01, \*\*\**p* < 0.001. See Figure S5 for lack of effect of IN inactivation on eEPSCs.

To assess the role of Sst^+^ and PV INs in BLA FFI, we stimulated LEC and recorded eEPSCs and eIPSCs in BLA PNs using Sst-hM4Di and PV-hM4Di mice before and after bath application of CNO (10 μM). To determine the effect of IN inactivation on FFI we quantified two measures: eIPSC amplitude and inhibition-excitation balance (I/E balance). We defined I/E balance as the eIPSC amplitude for each cell normalized to its eEPSC amplitude. We examined this measure in addition to eIPSC amplitude to control for potential differences across cells with regards to the number of excitatory inputs to BLA activated by LEC stimulation. Consistent with the disynaptic nature of LEC-driven FFI in BLA (Figure 1), LEC stimulation elicited an eIPSC that was significantly delayed relative to the eEPSC in BLA PNs in both Sst-hM4Di and PV-hM4Di mice in control conditions and in the presence of CNO (Figure 5B; Sst-hM4Di: latency_Sst-EPSC-Ctrl_ = 7.01 ± 0.26 ms, latency_Sst-IPSC-Ctrl_ = 8.57 ± 0.38 ms, latency_Sst-EPSC-CNO_ = 6.77 ± 0.26 ms, latency_Sst-IPSC-CNO_ = 8.57 ± 0.31 ms, *p*_Sst-Ctrl_ = 0.0018, *p*_Sst-CNO_ = 8.80 × 10^-4^, paired t-tests, n_Sst_ = 13, 6; PV-hM4Di: latency_PV-EPSC-Ctrl_ = 7.28 [5.89/7.41] ms, latency_PV-IPSC-Ctrl_ = 9.52 [7.40/10.30] ms, latency_PV-EPSC-CNO_ = 6.87 ± 0.27 ms, latency_PV-IPSC, CNO_ = 9.32 ± 0.66 ms, *p*_PV-Ctrl_ = 2.44 × 10^-4^, WSR test, *p*_PV-CNO_ = 0.0042, paired t-test, n_PV_ = 13, 7). Importantly, these data indicate that hM4Di expression or CNO application does not alter the ability of LEC stimulation to drive FFI. When we perfused CNO to inactivate the different IN subtypes, CNO reduced eIPSC amplitude and I/E balance in PNs from Sst-hM4Di mice by 30.2% and 40.2% respectively but had no effect on either measure in PNs from PV-hM4Di mice (Figure 5C, D; Sst-hM4D_i_: IPSC_Sst-Ctrl_ = 223.27 [131.10/439.99] pA, eIPSC_Sst-CNO_ = 155.87 [108.44/388.54] pA, I/E balance_Sst-Ctrl_ = 3.26 [1.85/4.88], I/E balance_Sst-CNO_ = 1.95 [1.36/2.58], *p*_Sst-IPSC_ = 0.040, *p*_Sst-IE_ = 0.013, WSR tests, n_Sst_ = 13, 6; PV-hM4D_i_: IPSC_PV-Ctrl_ = 238.06 ± 45.08 pA, eIPSC_PV-CNO_ = 201.87 ± 28.20 pA, I/E balance_PV-Ctrl_ = 3.53 ± 0.65, I/E balance_PV-CNO_ = 3.63 ± 0.67, *p*_PV-IPSC_ = 0.35, *p*_PV-IE_ = 0.85, paired t-tests, n = 13, 7). Finally, demonstrating the effects of Sst^+^ and PV^+^ IN inactivation were specific to FFI, bath application of CNO had no effect on eEPSCs in either mouse line (Figure S5).

To control for off-target effects of CNO (Gomez et al., 2017), we repeated the experiments in wildtype (WT) littermates of Sst-hM4Di and PV-hM4Di mice. In WT littermates, LEC stimulation elicited the delayed EPSC-IPSC pairing consistent with FFI (Figure 5b; latency_EPSC-Ctrl_: 6.63 [6.41/7.17] ms, latency_IPSC-Ctrl_: 9.27 [7.39/10.14] ms, latency_EPSC-CNO_: 6.75 ± 0.30 ms, latency_IPSC-CNO_: 8.50 ± 0.42 ms, *p*_Ctrl_ = 0.0024, WSR test, *p*_CNO_ = 0.0012, paired t-test, n= 12, 6), and CNO application had no effect on LEC-driven excitation or FFI in BLA (Figure 5e; Figure S5; IPSC_Ctrl_ = 172.29 [135.00/313.01] pA, IPSC_CNO_ = 189.68 [91.11/550.64] pA, I/E balance_Ctrl_ = 1.64 [0.83/7.30], I/E balance_CNO_ = 2.34 [1.31/5.13], *p*_IPSC_ = 0.73, *p*_IE_ = 0.68, WSR tests, n = 12, 6). Taken together with our data demonstrating that LEC afferents selectively synapse onto FS Sst^+^ INs among the Sst^+^ IN subtypes (Figures 2-3), these data show that FS Sst^+^, but not PV^+^ or nFS Sst^+^, INs mediate LEC-driven FFI in BLA.

### Sst^+^ INs control BLA plasticity

Because FS Sst^+^ INs mediate LEC evoked FFI (Figure 5), we hypothesized that Sst^+^, but not PV^+^, INs would gate plasticity at the LEC→BLA synapse. To test this hypothesis, we used the Hebbian-like pairing protocol that we showed in Figure 1 induces a bidirectional plasticity in BLA PNs in a manner gated by local GABAergic inhibition. We reasoned that if we inactivate the IN subtype that gates plasticity, the number of PNs responding to the pairing protocol with long-term plastic changes should increase. We conducted plasticity experiments with the Sst-hM4Di and PV-hM4Di mice and included 10 μM CNO in the bath to selectively inactivate either cell type. Inactivating Sst^+^ INs while stimulating PNs with the pairing protocol induced bidirectional plasticity in 6 of 7 neurons (Figure 6A; LTP: 141.79 ± 7.33 %, n = 3, 3; LTD: 66.28 ± 11.24 %, n = 3, 3). However, when we ran PNs through the pairing protocol while inactivating PV^+^ INs, the majority of PNs did not undergo plastic changes. Specifically, in 5 of 7 PNs there was no effect on eEPSC amplitude (Figure 6B; No plasticity PNs: 99.23 ± 4.20 %, n = 5, 4). Finally, we compared the proportion of PNs undergoing long-term plastic changes across conditions and found that a significantly greater proportion of neurons underwent plastic changes when we selectively inactivated Sst^+^ INs compared to PV^+^ INs (Figure 6C; Proportion plastic: Sst-hM4Di + CNO = 0.86, PV-hM4Di + CNO = 0.29; *p* = 0.031, X^2^ test, n_Sst_ = 7, 6, n_PV_ = 7, 6). Thus, we found that Sst^+^, but not PV^+^, INs gate plasticity at the LEC→BLA synapse.

**Figure 6.**
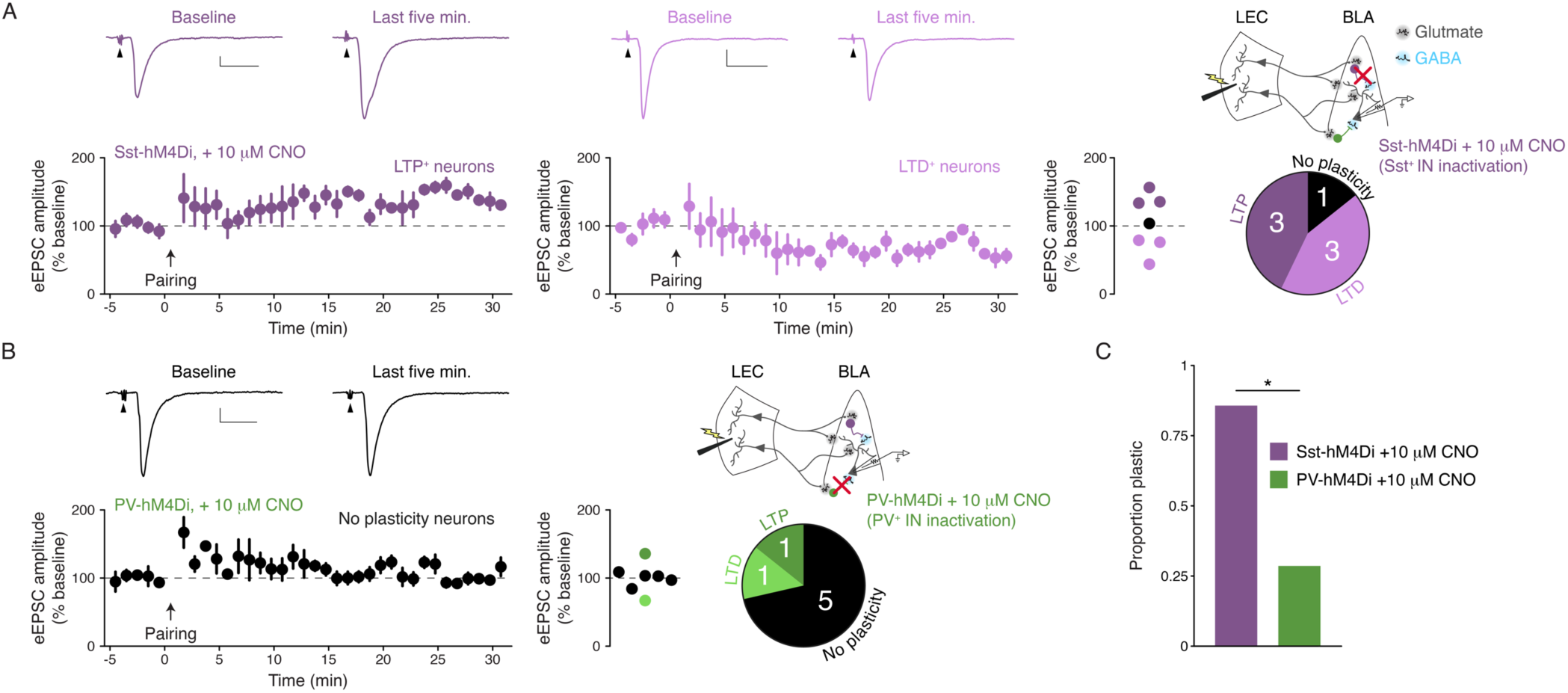
Sst^+^ INs control BLA plasticity. (A) Plasticity experiments in the Sst-hM4Di + 10 μM CNO condition. Sst^+^ IN inactivation during pairing of pre- and postsynaptic activity leads to LTP and LTD. Top: representative mean eEPSC trace for baseline period and last five minutes of experiment for neurons that underwent LTP (left; scale bars: 20 pA, 20 ms) or LTD (right; scale bars: 50 pA, 20ms). Arrowheads: truncated stimulus artifacts. Bottom: eEPSC (% baseline) amplitude as a function of time; data displayed as mean ± s.e.m. Right: Scatter plot of eEPSC amplitude (% baseline) in the last five minutes of the experiments (n = 7, 6). Pie chart shows results of experiments. (B) As for A, but in the PV-hM4Di + 10 μM CNO condition. Plasticity changes are rarely observed when PV^+^ IN inactivation is paired with the induction protocol (scale bars: 20 pA, 20 ms; n = 7, 6). (C) Significantly more cells undergo long-term plastic changes Sst^+^ IN inactivation compared to PV^+^ IN inactivation (X^2^ test: *p* = 0.031, n_Sst_ = 7, 6, n_PV_ = 7, 6). Data points are colored depending on results of experiment (*p* < *α*_FDR_): significant LTP experiments, significant LTD experiments, or no significant plasticity.

## DISCUSSION

Our data show that a previously unreported subpopulation of fast spiking Sst^+^ INs mediates LEC-driven FFI to gate plasticity in BLA (Figure 7). Although LEC afferents synapse onto both PV^+^ and FS Sst^+^ INs (Figures 2, 3), only FS Sst^+^ INs fired in response to LEC activity (Figure 4). Inactivation of Sst^+^, but not PV^+^, INs led to a reduction in FFI (Figure 5) and to an increase in plastic changes at LEC→BLA synapses that mirrored those seen during a complete GABA_A_ receptor blockade (Figures 1, 6). Our findings are consistent with prior experiments in BLA indicating that FFI gates plasticity onto PNs (Bazelot et al., 2015; Bissière et al., 2003; Tully et al., 2007) and that Sst^+^ INs form the anatomical circuit underlying FFI (Duvarci and Paré, 2014; Smith et al., 2000; Unal et al., 2014). In summary, these data illuminate a circuit mechanism for the feedforward inhibitory control mediated by entorhinal afferents onto BLA PNs (Lang and Paré, 1997), which suggests that this circuit is likely involved in olfactory and other forms of sensory-valence learning (Keene et al., 2016; Kitamura et al., 2017; McDonald, 1998; Mouly and Di Scala, 2006; Schoenbaum et al., 1999; Tsao et al., 2013; Xu and Wilson, 2012).

**Figure 7.**
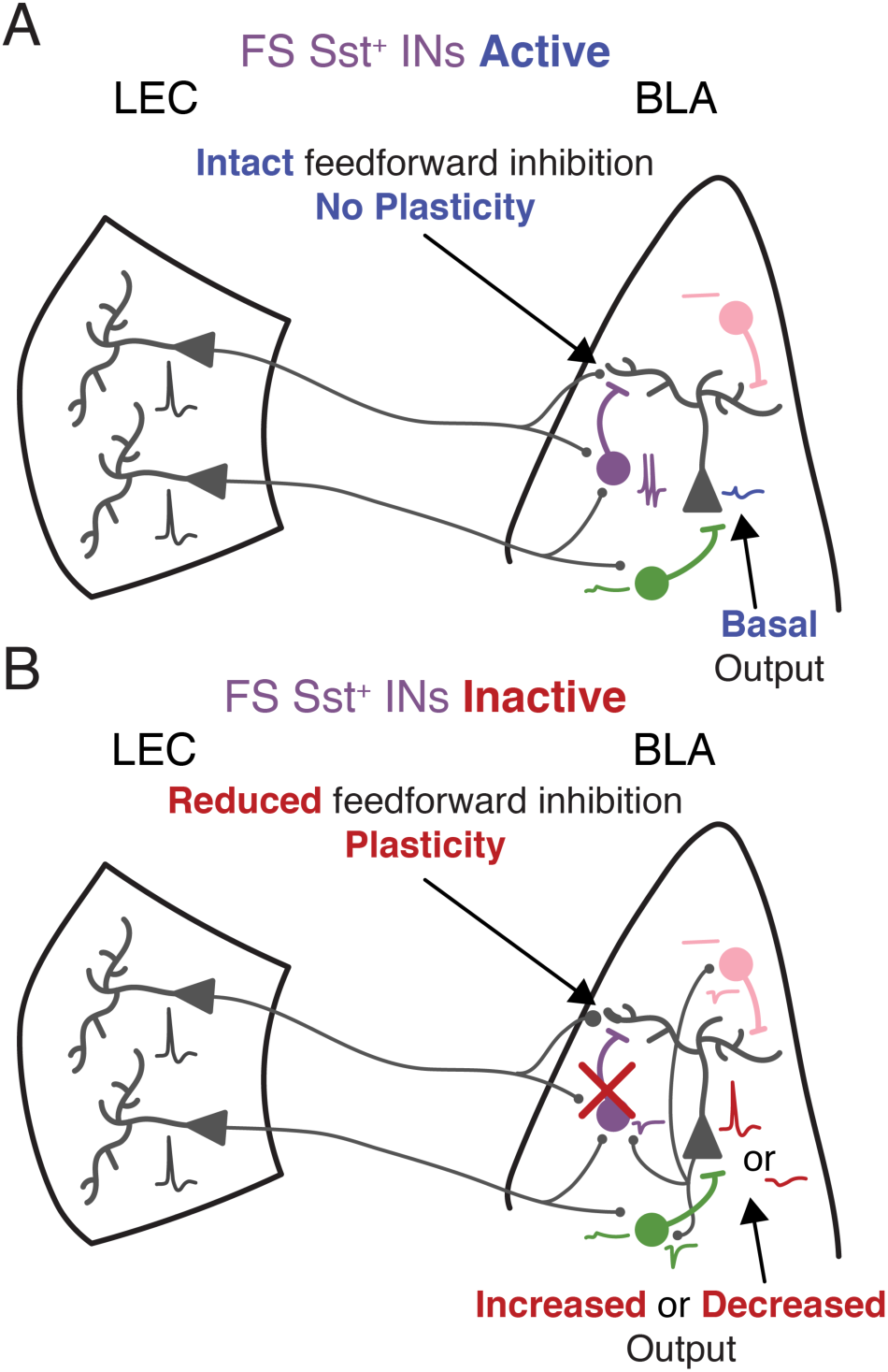
Schematic of findings. FS Sst^+^ INs (purple) fire APs in response to LEC input. nFS Sst^+^ INs (pink) do not respond to LEC input. PV^+^ INs (green) respond with a subthreshold excitatory response to LEC input. (A) When FS Sst^+^ IN mediated FFI is intact, BLA PNs will output their basal response to LEC input. (B) Inactivation of FS Sst^+^ IN removes the feedforward inhibitory control over BLA PNs, allowing them to undergo plastic changes. PV^+^ INs receive large uEPSC input from local BLA PNs compared to both Sst^+^ IN subtypes.

BLA Sst^+^ INs are a heterogenous population of dendritic targeting INs with diverse firing properties (Krabbe et al., 2018; Muller et al., 2007; Sosulina et al., 2010; Wolff et al., 2014). This is similar to what is observed in cerebral cortex (Tremblay et al., 2016). Accordingly, our cluster analysis and predictive modeling techniques revealed two distinct subpopulations of Sst^+^ INs that could be distinguished based on membrane properties, most notably maximum firing rate, hyperpolarization induced sag, and AP halfwidth (Figure 3). These membrane properties suggest differential expression of HCN and K_v_3 channels in the Sst^+^ IN subpopulations (Poolos et al., 2002; Rudy and McBain, 2001). Consistent with this, K_v_3.2 is expressed in a large subpopulation of BLA Sst^+^ INs (McDonald and Mascagni, 2006).

How IN heterogeneity maps onto specific functions within the BLA is critical for understanding survival circuits. We observed that our cluster analysis based on intrinsic properties separated the populations of Sst^+^ INs based on whether they displayed a synaptic response to LEC stimulation: 87.5% of Sst^+^ INs with a synaptic response were classified as fast spiking and 89.5% of Sst^+^ without a synaptic response were non-fast spiking. Although canonically PV^+^ INs are thought to be synonymous with FS INs, our data indicate that, at least in BLA, this is not the case. In BLA, our data reveal two functionally distinct populations of FS INs can be separated based on PV or Sst expression. Further, taken together with recent descriptions of FS Sst^+^ INs in cerebral cortex (Large et al., 2016; Ma et al., 2006; Nigro et al., 2018), we argue that a fast spiking phenotype is insufficient to classify an IN solely as PV^+^. Fast spiking Sst^+^ INs appear to be a biophysically and functionally distinct class of Sst^+^ INs that mediate feedforward inhibition to gate plasticity in the LEC→BLA circuit.

Whether the findings of the LEC→BLA circuit extend to other afferents to BLA is an important question (e.g. thalamic and other cortical inputs). BLA PV^+^ INs receive relatively little input from a variety of cortical sources (Smith et al., 2000). Instead, cortical inputs target PV^-^/calbindin^+^ INs (Unal et al., 2014). Interestingly, the major population of PV^-^/calbindin^+^ INs in BLA are Sst^+^ INs (McDonald and Mascagni, 2002). Additionally, FFI constrains plasticity at both thalamic (Bissière et al., 2003; Tully et al., 2007) and non-thalamic (Bazelot et al., 2015) inputs to BLA. Finally, the suppression of Sst^+^ IN activity is necessary for learning in an *in vivo* threat conditioning paradigm (Wolff et al., 2014). Thus, our findings will likely hold across a variety of inputs to the BLA.

Our data demonstrating that PV^+^ INs receive stronger uEPSCs from local PNs compared to Sst^+^ INs (Figure 4) and other recent work showing that PV^+^ INs fire APs more readily in response to local PN activation compared to cholecystokinin expressing INs (Andrási et al., 2017) are consistent with PV^+^ INs mediating feedback inhibition. A recent study (Lucas et al., 2016) shows that PV^+^ INs mediate polysynaptic inhibition in the lateral but not basal nucleus of the amygdala following stimulation of cortical (external capsule) or thalamic (internal capsule) fiber tracts that provide the input to the BLA. As stimulation of these fiber tracts leads to activation of local PNs and feedback inhibition (Szinyei et al., 2000), their results could be indicative of PV^+^ INs mediating BLA feedback inhibition, as has been suggested previously (Duvarci and Paré, 2014).

In contrast with the canonical BLA coronal slice, the horizontal slice prepation we used does not allow for unambigious discrimination between the lateral and basal nuclei; therefore, we cannot determine the subnucleus where we conducted our recordings. However, since the majority of LEC afferents target the basal nucleus (McDonald, 1998), it is conceivable that we recorded predominantly from neurons in the basal nucleus. If this were the case, this would provide an alternative explanation for the apparent discrepancy between our data and the data reported by Lucas and colleagues (Lucas et al., 2016). Nevertheless, FFI gates plasticity at a variety of different inputs to BLA (Bazelot et al., 2015; Bissière et al., 2003; Tully et al., 2007). Additionally, postsynaptic Ca^2+^ signalling plays a major role in amygdalar plasticity (Bauer et al., 2002; Humeau et al., 2005; Humeau and Lüthi, 2007; Weisskopf et al., 1999), and dendritic inhibition regulates postynaptic Ca^2+^ dynamics in pyramidal neuron dendrites (Chiu et al., 2013; Higley, 2014; Miles et al., 1996; Müllner et al., 2015). Finally, the role of Sst^+^ INs in gating BLA-dependent learning *in vivo* (Wolff et al., 2014) and the relative dearth of cortical afferents targeting PV^+^ INs in BLA (Smith et al., 2000) suggest that our data likely generalize across both subnuclei of the BLA and a variety of inputs to the structure. Future experiments will be necessary to test this premise.

The findings of the present study differ substantially from previous reports on feedforward inhibition in cerebral cortex and hippocampus. In these structures, which share similar neuronal subtypes with BLA (Duvarci and Paré, 2014), PV^+^, and not Sst^+^, INs mediate feedforward inhibition (Glickfeld and Scanziani, 2006; Tremblay et al., 2016). Why would this be the case? We believe that our finding that FS Sst^+^ INs are the circuit mechanism underlying BLA plasticity holds the key to answering this question. In cortex and hippocampus, FFI exerts control over the *temporal fidelity of AP output* of PNs (Gabernet et al., 2005; Pouille and Scanziani, 2001) whereas in BLA, FFI plays a critical role in exerting control over the *plasticity* of PNs (Bazelot et al., 2015; Bissière et al., 2003; Tully et al., 2007).

Consistent with the role of FFI in cortex and hippocampus, the major circuit role of the perisomatic PV^+^ INs in cerebral cortex, hippocampus, and BLA is to regulate the AP output of PNs (Glickfeld and Scanziani, 2006; Pouille and Scanziani, 2001; Tremblay et al., 2016; Woodruff and Sah, 2007b). Conversely, BLA Sst^+^ INs primarily target the dendrites of PNs (Muller et al., 2007; Wolff et al., 2014). The major circuit role of dendritic targeting INs is to regulate the local Ca^2+^ flux in dendrites (Chiu et al., 2013; Miles et al., 1996; Müllner et al., 2015). Further, recent work in cerebral cortex demonstrates that dendritic disinhibition via Sst^+^ INs inhibition is critical in gating cortical pyramidal neuron plasticity *ex vivo* and in gating learning *in vivo* via regulation of learning dependent changes in pyramidal neuron activity (Adler et al., 2019; Williams and Holtmaat, 2019). Additionally, our data demonstrate that PV^+^ INs receive stronger synaptic input from local BLA PNs compared to Sst^+^ INs (Figure 4). This finding compliments prior work showing that BLA PV^+^ INs are readily recruited by low levels of activity in local PNs (Andrási et al., 2017) and are connected via electrical synapses to synchronize their activity across groups of PV^+^ INs (Woodruff and Sah, 2007a). Thus, PV^+^ INs are positioned to provide feedback and lateral inhibition to control BLA output whereas FS Sst^+^ INs are positioned to regulate plasticity via feedforward inhibition. Together, the inhibitory networks would work in concert to fine-tune patterns of BLA activity to allow animals to learn about and react to stimuli. Our results illuminate an important point: archetypal circuit motifs that reappear across brain regions may be realized by different types of neurons in these brain areas depending on the functional role of the local circuit motif.

How might the BLA microcircuitry achieve the ability of BLA to appropriately gate sensory stimuli to drive goal directed behavior in a multi-sensory environment? We propose disinhibition of PN dendrites by the suppression of FS Sst^+^ IN activity as the microcircuit mechanism by which the BLA gates stimuli. Indeed, dendritic inhibition and disinhibition are known to play important roles in learning and behavior (Adler et al., 2019; Higley, 2014; Letzkus et al., 2015). A recent modeling study explored how these circuit mechanisms would be engaged and found increased disinhibition of pyramidal neuron dendrites improves the ability of a model cortical layer 2/3 circuit to discriminate between different sensory stimuli (Yang et al., 2016). Adding PV^+^ IN mediated perisomatic inhibition to the model circuit further improved its ability to selectively gate sensory input, and this circuit organization provided a framework for efficient Hebbian-like plasticity mechanisms (Yang et al., 2016). This pattern of activity is seen *in vivo* in BLA during threat conditioning: sensory stimuli drive increased PV^+^ IN firing and inhibit Sst^+^ IN firing (Wolff et al., 2014). Further decreasing Sst^+^ or increasing PV^+^ IN activity facilitated learning whereas increasing Sst^+^ or decreasing PV^+^ IN activity impaired learning (Wolff et al., 2014). Taken in the context of these findings, our data provide evidence for a circuit mechanism explaining how this pattern of activity *in vivo* may leads to learning: (1) the inhibition of Sst^+^ INs in response to a sensory stimulus would lead to dendritic disinhibition to select the specific sensory stimulus; and, (2) this would allow for plastic changes in the BLA PN to determine appropriate behavioral responses to the sensory stimulus in the future. The activation of PV^+^ INs could then organize the PNs to select the correct sensory-valence engrams, likely via feedback inhibition (Andrási et al., 2017; Duvarci and Paré, 2014). Importantly, other modeling studies have demonstrated that feedback inhibition plays a crucial role in the stability of BLA engrams (Feng et al., 2016).

Our data show that the inhibition of BLA Sst^+^ INs promotes the plasticity of cortical synapses onto BLA PNs (Figure 6). This is consistent with previous findings that BLA Sst^+^ INs must be inhibited for the duration of the sensory-valence pairing (Wolff et al., 2014). If the inactivation of Sst^+^ INs to decrease FFI is the circuit mechanism underlying the plasticity of BLA PNs, how does it occur *in vivo*? One possibility is inhibition from PV^+^ INs. However, this appears unlikely. Our data demonstrate that LEC afferents are inefficient at driving PV^+^ INs to spike, consistent with anatomical findings showing a paucity of cortical afferents targeting PV^+^ INs (Smith et al., 2000). Although sensory stimuli activate BLA PV^+^ INs (Wolff et al., 2014), our data and other studies suggest this is likely indirect and due to their activation by local PNs to mediate feedback inhibition (Andrási et al., 2017; Smith et al., 2000). Additionally, BLA PV^+^ INs are inhibited by the presentation of the valence signal (Wolff et al., 2014); thus, they cannot be the source of the inhibition of Sst^+^ INs during the sensory-valence pairing.

An alternative possibility for mediating the disinhibition of BLA PN dendrites is neuromodulation. Neuromodulation plays an important role in BLA dependent learning processes (Johansen et al., 2011). For instance, one candidate for mediating the inhibition of Sst^+^ INs *in vivo* is dopamine. Dopamine acts via the D2 dopamine receptor to reduce FFI in BLA and allow LTP to occur in *ex vivo* slice preparations (Bissière et al., 2003), reduces GABA release from BLA Sst^+^ INs (Chu et al., 2012), and regulates learning induced changes in BLA PN responses to sensory stimuli *in vivo* (Rosenkranz and Grace, 2002). A recent study demonstrated that the majority of dopaminergic afferents that target the BLA originate in the dorsal tegmental regions and that inactivation of these dorsal tegmental dopamine neurons during learning impaired later recall of sensory-valence associations (Groessl et al., 2018). Finally, it has been proposed that evolutionary pressures drive the selection of specific neural microcircuit mechanisms for fundamental, species-specific survival behaviors (Anderson and Adolphs, 2014; LeDoux, 2012). Consistent with this notion, in *Drosophila* dopamine release during learning acts via the *Drosophila* dopamine 2-like receptor to reduce GABAergic FFI onto Kenyon cells in the mushroom body (Zhou et al., 2019). This dopamine-induced reduction in FFI allows the Kenyon cells to undergo the necessary plasticity to learn sensory-threat associations (Zhou et al., 2019). Taken together, these data suggest that dopamine release in BLA *in vivo* could act on local Sst^+^ INs to reduce the sensory afferent driven FFI onto BLA PNs to allow for the plastic changes that underlie the sensory-valence learning to occur; however, future experiments are necessary to directly test this hypothesis. Given the degeneracy inherent to neural circuits and neuromodulation (Marder et al., 2014), other ways of inhibiting Sst^+^ INs *in vivo* are likely and warrant future study.

BLA plasticity has been implicated in olfactory learning (Schoenbaum et al., 1999) and the LEC→BLA synapse can undergo plastic changes *in vivo* (Yaniv et al., 2003). In addition, stimulation of LEC elicits heterosynaptic inhibition of responses to stimulation of the olfactory bulb in piriform cortex and BLA (Mouly and Di Scala, 2006), and optogenetic activation of BLA projection fibers modifies odorant responsiveness of neurons in posterior piriform cortex (Sadrian and Wilson, 2015). Finally, following learning induced plasticity, BLA PNs organize into engrams whose future activation is sufficient to drive the learned goal-directed behavior (Han et al., 2009; Hsiang et al., 2014; Redondo et al., 2014). BLA neurons begin to encode the valence associated with odors and other sensory stimuli within a few trials of sensory-valence pairing whereas it takes tens of trials for neurons in the olfactory system to develop valence coding (Doucette and Restrepo, 2008; Paton et al., 2006; Schoenbaum et al., 1999). Our findings raise the question whether the FS Sst^+^ IN mediated FFI is involved in changes in odorant processing that take place during olfactory associative learning (Doucette et al., 2011; Gire et al., 2013; Jordan et al., 2018). Specifically, we propose that following learning BLA PNs impress the odor-valence association onto the olfactory bulb and regions of olfactory cortex where olfactory associative learning induced plasticity occurs.

The BLA has been implicated in a variety of neuropsychiatric diseases ranging from anxiety to addiction (Duvarci and Paré, 2014; Janak and Tye, 2015). Our finding that FS Sst^+^ IN mediated FFI is the circuit mechanism gating BLA plasticity suggests that *in vivo* manipulation of this cell population may have therapeutic potential for conditions like anxiety where aberrant BLA-dependent learning is thought to be the underlying cause (Duvarci and Paré, 2014).

## Supporting information

Key Resources Table

## Acknowledgments

We would like to thank Nicole Arevalo and the University of Colorado Anschutz Medical Campus Breeding Core for their assistance with animal care. We would also like to thank members of the Huntsman and Restrepo labs and Alicia M. Purkey for discussions on the data. This work was supported by U.S. National Institutes of Health grants (R01 DC000566 to D.R., R01 NS095311 to M.M.H., T32 NS099042 to E.M.G., and T32 GM763540 to J.D.G.) and U.S. National Science Foundation Graduate Research Fellowship (DGE-1553798 to E.M.G.).

## The author contributions

Supervision, D.R. and M.M.H.; Conceptualization, E.M.G., D.R., and M.M.H; Electrophysiology investigation, E.M.G.; Immunohistochemistry investigation, J.D.G., and M.M.; Software, E.M.G., P.C., and S.M.B.; Formal Analysis, E.M.G, J.D.G., M.M., and P.C.; Visualization, E.M.G., J.D.G., and M.M.; Resources, K.R.S, D.R., and M.M.H.; Writing – Original Draft, E.M.G. Writing – Review & Editing, E.M.G., J.D.G., M.M., P.C., S.M.B., K.R.S., D.R., and M.M.H.; Funding Acquisition, E.M.G., J.D.G., D.R., and M.M.H.

## Methods

### Contact for Reagent and Resource Sharing

Further information and requests for resources and reagents should be directed to and will be fulfilled by the Lead Contact, Molly M. Huntsman.

### Experimental Model and Subject Details

All experiments were conducted in accordance with protocols approved by the Institutional Animal Care and Use Committee at the University of Colorado Anschutz Medical Campus. Slice electrophysiology experiments were conducted on mice aged postnatal days 35-70. Immunohistochemistry experiments were conducted on mice aged postnatal days 60-120. Experiments were conducted regardless of the observed external genitalia of the mice at weaning. To track estrous, vaginal swabs were collected from mice with vaginas that were used in experiments. The following mouse lines were used in the experiments: C57Bl/6J (Jackson Lab #000664), Sst-tdTomato, PV-tdTomato, Sst-hM4Di, PV-hM4Di, and wildtype littermates of the Sst-hM4Di and PV-hM4Di mice. Sst-tdTomato, PV-tdTomato, Sst-hM4Di, and PV-hM4Di mouse lines were generated by crosses of Sst-*ires*-Cre (Jackson Lab #013044) and PV-*ires*-Cre (Jackson Lab #008069) lines with either the Rosa-CAG-LSL-tdTomato-WPRE (*Ai9*; Jackson Lab #007905) or the R26-hM4Di/mCitrine (Jackson Lab #026219) lines. Please see Key Resources Table for more details on strain information.

### Method Details

#### Acute slice preparation for electrophysiology

Animals were first anesthetized with CO_2_ and decapitated. Brains were quickly dissected and placed in an ice-cold, oxygenated (95% O_2_-5% CO_2_) sucrose-based slicing solution (in mM: sucrose, 45; glucose, 25; NaCl, 85; KCl, 2.5; NaH_2_PO_4_, 1.25; NaHCO_3_, 25; CaCl_2_, 0.5; MgCl_2_, 7; osmolality, 290-300 mOsm/kg). 300-400µm horizontal slices were obtained using a vibratome (Leica Biosystems, Buffalo Grove, IL, USA). Slices were incubated in oxygenated (95% O_2_-5% CO_2_) artificial cerebral spinal fluid (ACSF; in mM: glucose, 10; NaCl, 124; KCl, 2.5; NaH_2_PO_4_, 1.25; NaHCO_3_, 25; CaCl_2_, 2; MgCl_2_, 2; osmolality 290-300 mOsm/kg) at 36°C for at least 30 minutes. All reagents were purchased from Sigma-Aldrich (St. Louis, MO, USA).

#### Electrophysiology

Slices were placed in a submerged slice chamber and perfused with ACSF heated to 32-37°C. Slices were visualized using a moving stage microscope (Scientifica: Uckfield, UK; Olympus: Tokyo, Japan) equipped with 4× (0.10 NA) and 40× (0.80 NA) objectives, differential interference contrast (DIC) optics, infrared illumination, LED illumination (CoolLED, Andover, UK), a CoolSNAP EZ camera (Photometrics, Tuscon, AZ, USA), and Micro-Manager 1.4 (Open Imaging, San Francisco, CA, USA). Whole cell patch clamp recordings were made using borosilicate glass pipettes (2.5-5.0 M*Ω*; King Precision Glass, Claremont, CA, USA) filled with intracellular recording solution. For voltage clamp experiments on FFI a cesium methanesulfonate (CsMe) based intracellular solution was used (in mM: CsMe, 120; HEPES, 10; EGTA, 0.5; NaCl, 8; Na-phosphocreatine, 10; QX-314, 1; MgATP, 4; Na_2_GTP, 0.4; pH to 7.3 with CsOH; osmolality adjusted to approximately 290 mOsm/kg). For all remaining voltage clamp experiments and for all current clamp experiments, a potassium gluconate based intracellular solution was used (in mM: potassium gluconate, 135; HEPES, 10; KCl, 20; EGTA, 0.1; MgATP, 2; Na_2_GTP, 0.3; pH to 7.3 with KOH; osmolality adjusted to approximately 295 mOsm/kg). A subset of the potassium gluconate recordings were supplemented with 0.2-0.5 % biocytin to allow for post-hoc morphological analysis. Access resistance was monitored throughout the experiments and data were discarded if access resistance exceeded 25 M*Ω* or varied by more than ± 20%. No junction potential compensation was performed. Data were acquired with a Multiclamp 700B amplifier and were converted to a digital signal with the Digidata 1440 digitizer using pCLAMP 10.6 software (Molecular Devices, Sunnyvale, CA). Data were sampled at 10 kHz and lowpass filtered at 4 kHz. Offline, current data were filtered using a 3^rd^ order Savistky-Golay filter with a ± 0.5 ms window after access resistance was assessed. Mean traces were created by first aligning all events by their point of maximal rise (postsynaptic currents) or by threshold (APs) and then obtaining the mean of all events; mean subthreshold EPSPs were not aligned prior to averaging.

#### Cell-type identification

##### Principal Neurons (PNs)

PNs were targeted based on their large, pyramidal-like soma. Recordings were terminated if the physiology of the neuron was inconsistent with BLA PNs (e.g. high membrane resistance, narrow AP halfwidth, large and fast spontaneous EPSCs).

##### Interneurons (INs)

INs were targeted based on fluorescence in the Sst-tdTomato, PV-tdTomato, SST-hM4D_i_, and PV-hM4D_i_ mouse lines. A 470 nm LED was used to identify mCitrine^+^ INs in SST-hM4D_i_ and PV-hM4D_i_ mouse lines, and a 535 nm LED was used to identify tdTomato^+^ INs in the SST-tdTomato and PV-tdTomato mouse lines (CoolLED, Andover, UK).

#### Pharmacology

DNQX, D-APV, and gbz were purchased from Tocris Biosciences (Bristol, UK) and CNO was purchased from Enzo Life Sciences (Farmingdale, NY). DNQX stock was made at 40 mM and diluted to a final concentration of 20 μM in ACSF; D-APV stock was made at 50 mM and diluted to a final concentration of 50 μM in ACSF; GBZ stock was made at 25 mM and diluted to a final concentration of 5 μM in ACSF; and, CNO stock was made at 10 mM and diluted to a final concentration of 10 μM in ACSF. All stocks were stored at −20°C and CNO was used within one month of making the stock solution.

#### Electrophysiology experimental design

##### FFI, voltage clamp

The LEC was stimulated using a bipolar stimulating electrode (FHC, Inc., Bowdoin, ME, USA). eEPSCs (V_hold_ = −70 mV) and eIPSCs (V_hold_ = 0 mV) were recorded from BLA PNs in response to LEC stimulation. To assess the effects of different drugs on the eEPSCs and eIPSCs, ACSF containing DNQX, D-APV, gbz, and/or CNO was perfused onto the slice for five minutes prior to and continuously during the experiment. Effects of DNQX/APV and gbz on eEPSCs and eIPSCs were recorded using in an unpaired design where some PNs were recorded under control conditions and in the presence of DNQX/APV and gbz (given sequentially with time for washout) whereas others were recorded under control conditions in the presence of DNQX/APV or gbz. Effects of CNO on eEPSCs and eIPSCs were examined with a paired design where all PNs were recorded in both control conditions and in the presence of CNO.

##### FFI, current clamp

Membrane voltage of BLA PNs was recorded in response to 5 stimulations of the LEC at 20 Hz. I_hold_ was adjusted such that V_rest_ of the PNs was approximately −60 mV. To assess the role of GABA_A_ receptor mediated inhibition on PN AP firing, gbz was perfused onto the slice for five minutes prior to and continuously during the experiment.

##### Current injections

Membrane voltage of BLA neurons was recorded in current clamp in response to a series of square hyperpolarizing and depolarizing current injections. Prior to initiation of the series of current injections, V_m_ of the BLA neurons was adjusted to approximately −60 mV. Each cell was subjected to two series of 600 ms square current injections: −100 pA to +100 pA at 10 pA intervals and −200 pA to +400 pA at 25 pA intervals. The data collected in these experiments were used to determine active and passive membrane properties of the neurons.

##### Minimal stimulation

eEPSCs (V_hold_ = −70 mV) were recorded in voltage clamp in BLA PNs, Sst^+^ INs, and PV^+^ INs in response to LEC stimulation. Stimulation intensity was adjusted such that LEC stimulation resulted in recorded eEPSCs having a success rate of approximately 50% and an all-or-none amplitude response.

##### Recruitment of BLA INs by LEC afferents

The median stimulation intensity necessary to observe putative uEPSCs in BLA PNs (273 μA × ms) was used as the empirically derived PN threshold stimulation intensity. Membrane voltage responses of BLA Sst^+^ and PV^+^ INs were recorded in current clamp in response to 5 stimulations of the LEC at 20 Hz at the empirically derived PN threshold stimulation intensity.

##### Paired PN-IN recordings

Paired recordings were made between BLA PNs and nearby Sst^+^ or PV^+^ INs. We used a 2.5 nA, 2 ms current injection to drive a single AP in the PN (I_hold_ adjusted such that V_m_ ≈ −60 mV. BLA PN APs were repeated at 0.25 Hz and the response of the IN (V_hold_ = −70 mV) was recorded.

##### Spontaneous EPSC recordings

sEPSCs were recorded for 5 minutes in INs (V_hold_ = −70 mV) with no drugs in the bath.

##### Pharmacological effects of CNO on membrane potential

Membrane voltage of mCitrine^+^ neurons in BLA was recorded in the presence of 20 μM DNQX, 50 μM D-APV, and 5 μM GBZ. To assess the effects of CNO on membrane potential, baseline V_m_ was allowed to stabilize and was recorded for 3 minutes in the absence of CNO. Following recording of baseline V_m_, ACSF containing 10 μM CNO was perfused onto the slice and V_m_ was recorded for an additional 10 minutes. V_m_ was separated into 30 second bins. *Δ*V_m_ was defined as the difference in V_m_ between the mean V_m_ during the 3 minutes of baseline recordings and the mean V_m_ during the last 3 minutes of CNO application.

##### IN control of BLA plasticity

All experiments were done in acute slices from wildtype mice with no drugs or 5 μM gbz in the bath or in acute slices from Sst-hM4Di/PV-hM4Di mice with 10 μM CNO in the bath. eEPSCs were recorded (V_hold_ = −70 mV) in BLA PNs in response to 0.067 Hz stimulation of LEC (4 stimuli/minute). After a 5 minute baseline of eEPSCs was recorded, plasticity was induced (in current clamp, I_hold_ adjusted such that V_m_ was set to approximately −70 mV) by pairing three APs (2 nA, 5 ms current injections) with three monosynaptic EPSPs elicited by LEC stimulation at 33.3 Hz. LEC stimulation was timed such the EPSP was recorded approximately 5-10 ms prior to AP induction. Immediately following each AP, V_m_ was depolarized to > −40 mV for 25ms. This induction protocol was repeated 15 × at 0.2 Hz. The plasticity induction protocol was run only if the baseline recordings were recorded within approximately 10 minutes of achieving whole-cell configuration; if the baseline recorded concluded after 10 minutes of achieving whole-cell, the experiments were terminated. Following plasticity induction, eEPSCs were recorded in response to LEC stimulation at 0.067 Hz for 30 minutes. Experiments were discarded if eEPSC amplitude did not achieve a stable level post-induction (significant deviation from the middle 5 minutes compared to the last 5 minutes of the post-induction period as determined by an unpaired t-test or a MWU test depending on the normality of the data). This was done to ensure that changes in eEPSC amplitude were related to plasticity induction and were not a confound of other phenomena. Our plasticity induction protocol was based on prior studies examining plasticity at cortical inputs to BLA (Humeau et al., 2005; Humeau and Lüthi, 2007). To determine if plasticity occurred, we compared eEPSC amplitude in the 5 minutes prior to induction to eEPSC amplitude in the last 5 minutes of the post-induction period. To correct for multiple comparisons (i.e. testing for a change in eEPSC amplitude across every cell in each genotype), we used the FDR method (Curran-Everett, 2000) to adjust the critical significance value from 0.05 to a FDR-corrected value. Because we were correcting for multiple comparisons, we used a unpaired t-test or MWU test (where appropriate), with an FDR adjusted *α* value for each condition, to determine if the change in eEPSC amplitude from baseline to the last 5 minutes of the post-induction period was significant. For wildtype mice with no drugs, *α*_FDR_ was set to 0.013; for wildtype mice with 5 μM gbz, *α*_FDR_ was set to 0.044; for Sst-hM4Di mice with 10 μM CNO, *α*_FDR_ was set to 0.043; for PV-hM4Di mice with 10 μM CNO, *α*_FDR_ was set to 0.014. We defined cells as undergoing LTP if they had a significant increase in eEPSC amplitude from baseline to the last 5 minutes of the post-induction period; we defined cells as undergoing LTD if they had a significant decrease in eEPSC amplitude from baseline to the last 5 minutes of the post-induction period. For display purposes, eEPSC amplitude was normalized to the mean baseline eEPSC amplitude, and eEPSCs were binned by the minute.

#### Definitions of electrophysiological parameters

##### eEPSC/eIPSC detection and amplitude

eEPSCs (V_hold_ = −70 mV) were defined as negatively deflecting postsynaptic events that exceeded the mean baseline current (500 ms before stimulation) by 6× the median absolute deviation of the baseline current and that occurred within 20 ms of the end of the electrical stimulus artifact. eIPSCs (V_hold_ = 0 mV) were defined as positively deflecting postsynaptic events that exceeded the mean baseline current by 6× the median absolute deviation of the baseline current and that occurred within 20 ms of the end of the electrical stimulus artifact. To ensure that the detected eEPSCs/eIPSCs were related to the stimulus, we subsampled 25% of the sweeps in the experiment (or 5 sweeps if the experiment consisted of <20 sweeps) and found the maximal peak negative (eEPSC detection) or positive (eIPSC detection) deflection from baseline in the 20 ms after the stimulus artifact in those sweeps. Then, we found the median peak time for those sweeps and repeated the analysis over all sweeps in the experiment with a detection threshold of 6× the median absolute deviation of the baseline current and with a window set to ± 5 ms (eEPSC detection) or ± 7.5 ms (eIPSC detection) around the median peak time. The amplitude of each eEPSC and eIPSC was defined as the difference between the peak amplitude of the detected eEPSC or eIPSC and the mean baseline current for that sweep. The eEPSC or eIPSC amplitude for each cell in an experiment was defined as the mean of the amplitudes recorded from that cell (eEPSC/eIPSC successes only; failures were not included in eEPSC/eIPSC amplitude calculation). Where eEPSC failure amplitude is reported on a per sweep basis, it was defined as the maximal negative deflection from the current trace within ± 5 ms of the mean current peak time for that cell. If all sweeps in an experiment were eEPSC/eIPSC failures, the mean eEPSC/eIPSC amplitude was defined as the maximal negative (eEPSC) or positive (eIPSC) deflection in the mean current trace that occurred within 20 ms of the end of the electrical stimulus artifact.

##### Success rate

Success rate was defined as 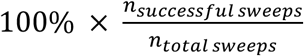 where successful sweeps were defined as sweeps where an eEPSC was detected.

##### I/E balance

I/E balance was defined as the ratio of eIPSC to eEPSC amplitude recorded in the same neuron.

##### EPSC/IPSC 20%-80% risetime

20%-80% risetime was defined as the time it took an EPSC or IPSC to reach 80% of its peak amplitude from 20% of its peak amplitude. 20%-80% risetime was calculated for each sweep unless obscured by a spontaneous event. The risetime for each cell was defined as the mean of all risetimes recorded from that cell.

##### EPSC/IPSC latency and jitter

Latency of EPSCs and IPSCs was defined as the time between the end of the electrical stimulation or the peak of the PN action potential (in paired recordings) and the point of 20% rise for an EPSC or IPSC as calculated for the 20%-80% risetime. The EPSC or IPSC latency for each cell was defined as the mean of the latencies recorded from that cell. The EPSC or IPSC jitter for each cell was defined as the standard deviation of the latencies recorded from that cell.

##### EPSC/IPSC τ_Decay_

EPSC *τ*_Decay_ was determined using a single exponential fit, 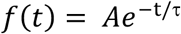. IPSC *τ*_Decay_ was defined as the weighted time-constant of IPSC decay. Briefly, a double exponential fit, 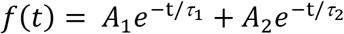, was used to obtain the parameters to determine the weighted time-constant where 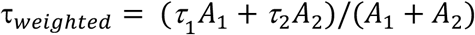. *τ*_Decay_ was calculated using the mean EPSC or IPSC trace for a cell.

##### uEPSC detection and amplitude

uEPSCs were defined as negative current deflections recorded in the IN that exceeded a detection threshold of 6× the median absolute deviation of the baseline current and occurred within 3ms of the PN AP peak during paired recordings. The amplitude of each uEPSC was defined as the difference between the peak amplitude of the detected uEPSC and the mean baseline current for that sweep. The uEPSC amplitude for each cell in an experiment was defined as the mean of the amplitudes recorded from that cell (successes only).

##### sEPSC detection and amplitude

sEPSCs were detected by a combined template and threshold method. Briefly, a template was made by subsampling 10% of local negative peaks exceeding at least 5 × the median absolute deviation of a rolling baseline current (50ms prior to the peak). The template current was then truncated from its 20% rise point through the end of the decay time constant for the template current. Next, all local negative peaks exceeding 5 × the median absolute deviation of a rolling baseline current (50ms prior to the peak) were collected. The template current was then scaled to each individual putative sEPSC peak and each sEPSC peak was assigned a normalized charge integral relative to the template. Finally, a normalized charge integral cutoff was chosen to exclude obvious noise/non-physiological events below a certain normalized charge integral. sEPSC amplitude was defined as the difference between the peak amplitude of each detected current and its corresponding baseline current. sEPSC for each cell was defined as the median peak amplitude for that cell.

##### sEPSC frequency

sEPSC frequency for each sESPC was defined as the inverse of the interevent intervals of the sEPSCs. The frequency measure for each neuron was defined as the median of the sEPSC frequencies for that cell.

##### Membrane resistance

Membrane resistance was defined as the slope of the best fit line of the I-V plot using the −100 pA to +100 pA (10 pA steps) series of current injections. Mean voltage response to each current injection step was defined as the difference between baseline mean membrane voltage (100 ms prior to current injection) and the mean membrane voltage during the 100 ms period from 50 ms after the start of the injection to 150ms after the start of the current injection. This 100 ms window was chosen to allow for measurement of the change in V_m_ after the membrane had charged and prior to any potential HCN channel activation. The I-V plot was constructed using all current steps below rheobase.

##### Maximum firing rate

Maximum firing rate was defined as the inverse of the inter-spike interval (ISI) during the first 200 ms of the most depolarizing current injection step before attenuation of AP firing was observed. Max FR was calculated using the −200 pA to +400 pA (25 pA steps) series of current injections.

##### AP threshold

AP threshold was defined as the voltage at which 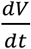 exceeded 20 V/s. AP threshold was calculated at the rheobase sweep of the −200 pA to +400 pA (25 pA steps) series of current injections.

##### AP amplitude

Amplitude of the AP was defined as the voltage difference between the peak of the AP and its threshold potential. AP amplitude was calculated at the rheobase sweep of the −200 pA to +400 pA (25 pA steps) series of current injections.

##### AP halfwidth

AP halfwidth was defined as the time between the half-amplitude point on the upslope of the AP waveform to the half-amplitude point on the downslope of the AP waveform. AP halfwidth was calculated at the rheobase sweep of the −200 pA to +400 pA (25 pA steps) series of current injections.

##### After-hyperpolarization potential (AHP) magnitude

AHP magnitude was defined as the difference between the most hyperpolarized membrane voltage of the AHP (occurring within 100 ms after AP threshold) and AP threshold. AHP magnitude and latency data were calculated at the rheobase sweep of the −200 pA to +400 pA (25 pA steps) series of current injections. *Δ*AHP data were calculated at the rheobase + 50 pA sweep of the −200 pA to +400 pA (25 pA steps) series of current injections.

##### AHP latency

AHP latency was defined as the time from AP threshold and the peak of the AHP. *ΔAHP. Δ*AHP was defined as the difference between the first and last AHP (*ΔAHP* = *AHP_last_* – *AHP_first_*).

##### AP phase plot

The AP phase plot was obtained by plotting the rate of change of the mean AP for each cell from the rheobase sweep of the −200 pA to +400 pA (25 pA steps) series of current injections as a function of the corresponding membrane voltage.

##### Latency to first AP

AP latency was defined as the time from the initiation of the current injection to the peak of the first AP. AP latency was calculated at the rheobase sweep of the −200 pA to +400 pA (25 pA steps) series of current injections.

##### Firing rate adaptation ratio (FR adaptation)

Firing rate adaptation was defined as the ratio of the first and the average of the last two ISIs, such that 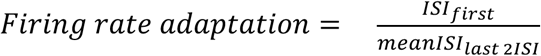. Firing rate adaptation was calculated at the rheobase +50 pA sweep of the −200 pA to +400 pA (25 pA steps) series of current injections.

##### AP broadening

AP broadening was defined as the ratio of the AP halfwidths of the first two APs 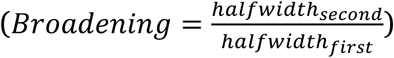. AP broadening was calculated at the rheobase +50 pA sweep of the −200 pA to +400 pA (25 pA steps) series of current injections.

##### AP amplitude adaptation

AP amplitude adaptation was defined as the ratio of the AP amplitude of the average of the last three APs and the first AP, such that 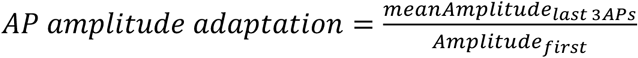. AP amplitude adaptation was calculated at the rheobase +50 pA sweep of the −200 pA to +400 pA (25 pA steps) series of current injections.

##### Membrane decay τ

Membrane decay *τ* was determined by using a single exponential fit, 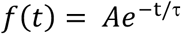, to fit the change in V_m_ induced by a −100 pA sweep in the −100 pA to +100 pA (25 pA steps) series of current injections.

##### Hyperpolarization-induced sag

Hyperpolarization-induced sag was calculated using the equation, 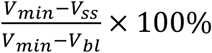, where V_min_ was defined as the most hyperpolarized membrane voltage during the current injection, V_ss_ was defined as the mean steady-state membrane voltage (last 200 ms of the current injection), and V_bl_ was defined as the mean baseline membrane voltage (100 ms prior to current injection). Hyperpolarization-induced sag was measured from the −200 pA current injection.

##### Rebound spikes

Rebound spikes were defined as the number of APs in the 500 ms following the −200 pA current injection.

##### APs per stimulus

The number of APs per stimulus was defined as the number of APs occurring within 50 ms of the stimulus. *V_rest_.* V_rest_ was defined as V_m_ (I_hold_ = 0 pA) during a 500 ms baseline prior to LEC stimulation during the experiments testing the recruitment of BLA INs by LEC afferents.

##### Subthreshold EPSP amplitude

Subthreshold EPSP amplitude was defined as the maximal, non-stimulus artifact, voltage deflection within 40 ms after LEC stimulation.

#### Immunohistochemistry

##### Biocytin filled neurons

To perform immunostaining of biocytin filled neurons, slices containing biocytin filled neurons were fixed in 4% PFA overnight at 4°C. After fixation, slices were transferred to PBS. Biocytin filled INs (n = 8 Sst^+^ Group 1; 3 Sst^+^ Group 2; 6 PV^+^) were blocked (1X PBS, 0.3% triton, 5% BSA, 5% Normal Donkey Serum) for 4 hours before 24-hour incubation with streptavidin conjugated Alexa Fluor 488 (1:500, ThermoFisher Scientific, Waltham, MA, USA) at 4°C. Slices were mounted with Prolong Gold and sealed for long-term storage. Slices were imaged using an Axio Observer microscope (Carl Zeiss, Okerkochen, Germany); equipped with a CSU-X1 spinning disc unit (Yokogawa, Musashino, Tokyo, Japan); 488 nm/40 mW laser; Plan-NeoFluar 40X (0.75 NA) air objective lens; and Evolve 512 EM-CCD camera (Photometrics, Tucson, AZ, USA). SlideBook 6.0 software (3i, Denver, CO, USA) enabled instrument control and data acquisition. Images were acquired in sections by following branched points from the cell soma. Images were stitched in Fiji software using Grid/Collection stitching (Preibisch et al., 2009) with an unknown position type.

##### Histological validation of PV^+^ and Sst^+^ IN identity

To perform the PV and Sst immunostaining, mice (n = 3 Sst-tdTomato mice, 3 PV-tdTomato mice) were sacrificed and transcardially perfused with ice cold 4% PFA (with 1.5% picric acid and 0.05% glutaraldehyde) followed by 30% sucrose protection. After the brain sank, coronal BLA slices of 30 μm thickness were obtained. Before application of blocking solution, slices were incubated for 30 minutes at room temperature with iFX-enhancer (Invitrogen, ThermoFisher Scientific, Waltham, MA, USA). After blocking, the slices were incubated with either rat anti-Sst antibody (1:100, MAB354, Millipore, Burlington, MA, USA) or guinea pig anti-PV antibody (1:500, 195004, Synaptic Systems, Göttingen, Germany) for at least 48 hours at 4°C. Then, a secondary antibody of either donkey anti-rat or donkey anti-guinea pig Alexa Fluor 488 (1:500, Jackson ImmunoResearch) was applied overnight at 4°C. The slices were imaged using a confocal laser scanning microscope (TCS SP5II, Leica Application Suite, Leica Biosystems, Buffalo Grove, IL, USA) with 10× 0.40 NA and 20× 0.70 NA dry objectives to determine the neuron identity.

#### Morphological analysis

##### Analysis of somatic and dendritic morphology

To determine morphological characteristics of biocytin filled neurons, stitched images were imported to Neurolucida (MBF Bioscience, Williston, VT, USA) to perform tracing. All analysis including sholl analysis (50 μM rings), dendrite branching, etc. was performed from traces using Neurolucida Explorer (MBF Bioscience).

### Quantification and Statistical Analysis

#### Statistical analyses

All data analysis (except decision tree and random forest analyses) were performed offline using custom written MATLAB code. Normality of the data were assessed using the Anderson-Darling test. For assessment of whether a single group differed from a normal distribution centered around zero, a one-sample t-test was used. For a test between two groups, a paired or unpaired t-test was used where appropriate. For tests between two groups of non-normal data, a Mann-Whitney U or Wilcoxon signed-rank test was used where appropriate. For tests between three or more groups of normal data with one independent variable, a one-way ANOVA was used with Tukey’s post-hoc test to examine differences between groups. A Kruskal-Wallis test was used to examine differences between three or more groups of non-normal data with one independent variable. A Mann-Whitney U test was used as a post-hoc test following a significant result in a Kruskal-Wallis test and was corrected for multiple comparisons using the FDR method (Curran-Everett, 2000). The critical significance value was set to *α* = 0.05 or was set to a FDR-corrected value (*α*_FDR_) for multiple comparisons. All statistical tests were two-tailed. Unless otherwise stated, experimental numbers are reported as *n* = x, y where x is the number of neurons and y is the number of mice. Statistical parameters are reported in the Results section and figure legends display *p* values and sample sizes.

#### Unsupervised cluster analysis

Unsupervised cluster analysis using Ward’s method (Ward, 1963) was used to classify Sst^+^ INs. Briefly, this method involves plotting each neuron in multidimensional space where each dimension corresponds to a given parameter. For our data, we plotted each Sst^+^ IN in 15-dimensional space (where each dimension corresponds to a z-score transformation of one of the 15 membrane properties obtained from the current injection experiments; values were z-score transformed so that parameters with large values, e.g. membrane resistance, would not influence the cluster analysis more than those with small values, e.g. halfwidth). From here, the analysis proceeded along *n* – 1 stages where *n* is the number of Sst^+^ INs. At stage *n* = 1, the two closest cells in the 15-dimensional space are grouped together. At subsequent stages, the closest cells are grouped together until only one group of all objects remains. We determined the final number of clusters by using the Thorndike procedure (Thorndike, 1953) where large distances between group centroids at one cluster stage relative to other stages are indicative of significant differences between groups (see Figure 3A, inset).

#### Decision tree analysis

Recursive partitioning analyses (Breiman et al., 1984) were conducted in R (1.01.136). The R package rpart (Therneau and Atkinson, n.d.) was used for recursive partitioning for classification. Decision trees were plotted using rpart.plot package, and random seeds were set using the rattle package. Predicted classes were determined from the unsupervised cluster analysis and were used to determine important discriminating parameters. The model was internally cross validated with a nested set of subtrees and the final model was selected as the subtree with the least misclassification error. Individual decision trees can suffer from overfitting. Therefore, the input parameters of the model were independently cross validated by bootstrapping 500 subsamples (without replacement and using random seeds). Each tree was pruned by choosing the complexity parameter with the lowest cross validated error. The mean correct prediction of the test set classifications (n = 20% of sample) by the pruned tree generated by modeling the training sets (n = 80% of sample) was 82.9%.

#### Random forest analysis

Supervised classification random forest was employed to reduce potential model overfitting. Random forest classification was conducted using the randomForest package in R (Liaw and Wiener, 2002). Random forest classification employs random subsampling of both parameters and bootstrapped subsamples of the dataset (with replacement). 10,000 decision trees were generated, and the modal tree was used as the final classification model. The out of bag estimate of the error rate of the model was 11.43%. An additional cross validation step was conducted by bootstrapping 80% of the initial sample to create the random forest model and tested against a subsample (n = 20% of sample; bootstrapped without replacement). The mean accuracy after 20 random forest runs was 90.7%. Gini impurity was calculated to determine parameter importance. Visualization of the frequency of individual samples falling within the same node across one run of the classification algorithm was obtained by calculating a proximity matrix for the samples and plotting it against the first two principle components.

#### Data display

Data visualizations were created in MATLAB and Adobe Illustrator. After analysis was completed, Neurolucida traces of soma and dendrites were thickened by 7 pixels (Figure 3) or 3-7 pixels (Figure S4) in Adobe Photoshop to improve visibility in figures. Normal data are presented as the mean ± s.e.m. Non-normal data are presented as the median with error bars extending along the interquartile range.

### Data and Software Availability

Data are available on request, and code is available on GitHub (https://github.com/emguthman/Manuscript-Codes).

