## Supplemental figures and table for "Cell type specific control of basolateral amygdala plasticity via entorhinal cortex-driven feedforward inhibition"

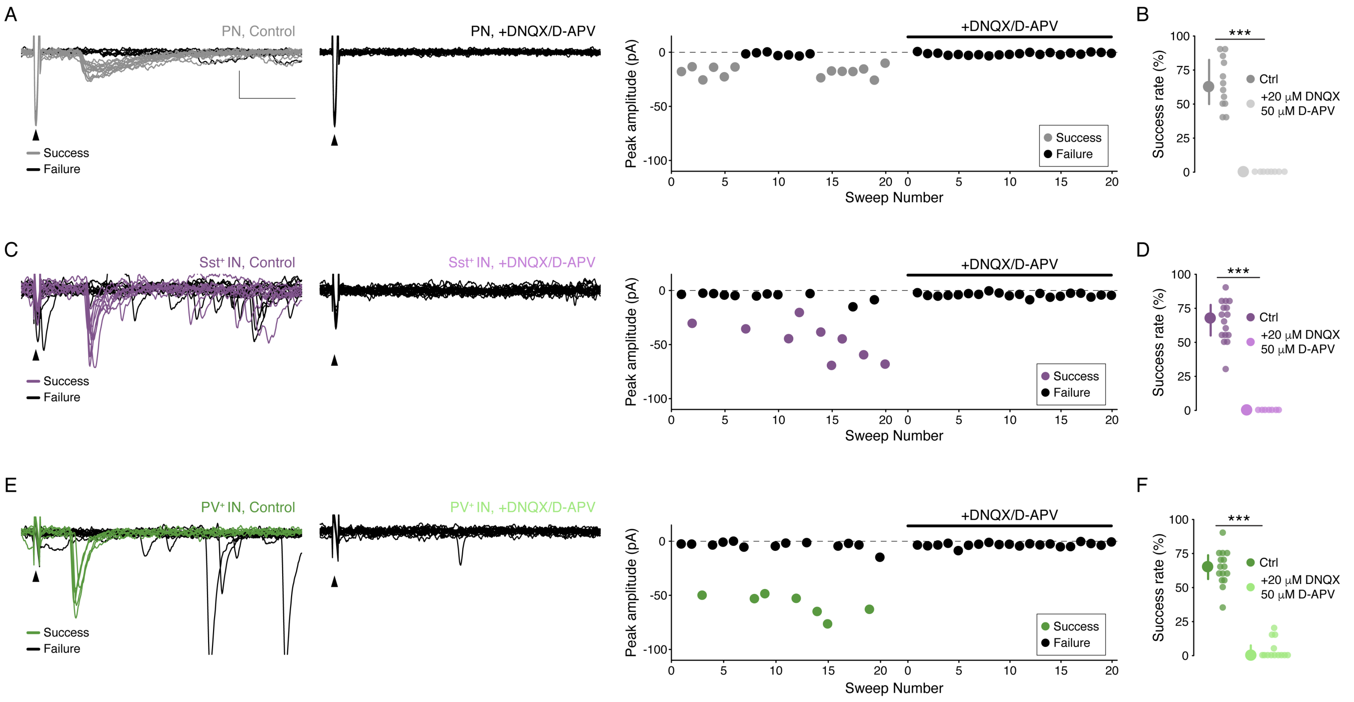


**Figure S1. Representative minimal stimulation experiments and glutamatergic nature of responses to LEC afferents. Related to Figure 2.**

(A) Minimal stimulation experiment in a representative PN. Left: 20 overlaid sweeps in control conditions. Middle: 20 overlaid sweeps in the presence of 20 µM DNQX and 50 µM D-APV. Right: Trial by trial summary of peak amplitudes. Arrowheads denote stimulation artifacts. Scale bars: 25 pA, 10 ms.

(B) Glutamate receptor blockade abolishes postsynaptic response in PNs (success $\text{rate}_{\text{Ctrl}}$ = 62.50 [50.00/82.50] %, success $\text{rate}_{\text{DNQX/APV}}$ = 0.00 [0.00/0.00] %, $p = 1.50\times10^{-4}$, MWU test; $n_{\text{Ctrl}}$ = 12 cells, 8 mice, $n_{\text{DNQX/APV}}$ = 8 cells, 7 mice; median [1$^{\text{st}}$ quartile/3$^{\text{rd}}$ quartile]).

(C) As (A), but for a representative Sst$^+$ IN.

(D) As (B), but for Sst$^+$ INs (success $\text{rate}_{\text{SCtrl}}$ = 67.5 [55.00/77.50] %, success $\text{rate}_{\text{DNQX/APV}}$ = 0.00 [0.00/0.00] %, $p = 1.62\times10^{-4}$, MWU test; $n_{\text{Ctrl}}$ = 16 cells, 8 mice; $n_{\text{DNQX/APV}}$ = 7 cells, 6 mice).

(E) As (A), but for a representative PV$^+$ IN.

(F) As (B), but for PV$^+$ INs (success $\text{rate}_{\text{Ctrl}}$: 65 [56.25/73.75] %, success $\text{rate}_{\text{DNQX/APV}}$ = 0.00 [0.00/7.50] %, $p = 5.33\times10^{-6}$, MWU test, $n_{\text{Ctrl}}$ = 15 cells, 5 mice, $n_{\text{DNQX/APV}}$ = 13 cells, 5 mice.

In panels B, D, and F, data shown as median with IQR. ***$p < 0.001$

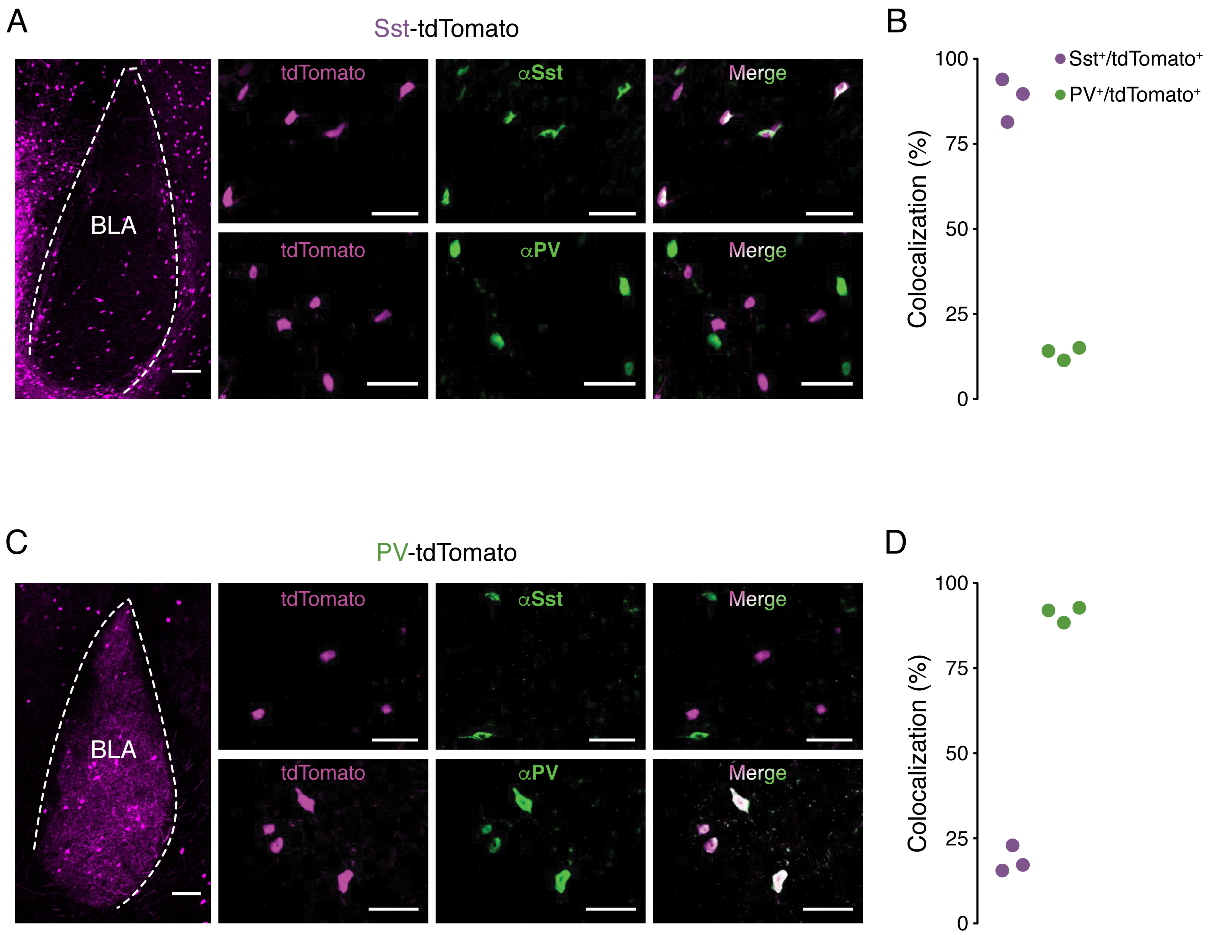


**Figure S2. Validation of PV-tdTomato and Sst-tdTomato mouse lines. Related to Figure 2.**

(A) Left: Representative confocal image of tdTomato fluorescence in a coronal section from a Sst-tdTomato mouse containing BLA. Scale bar: 100 μm. Middle: Representative confocal images of colocalization experiment in a Sst-tdTomato mouse. Left panel, tdTomato fluorescence. Middle panel, αSst or αPV immunofluorescence. Right panel, merge. Scale bar: 50 μm.

(B) Percent colocalization of αSst or αPV immunofluorescence with tdTomato fluorescence in Sst-tdTomato mice (88.09 ± 3.67% colocalization of αSst/tdTomato, 13.28 ± 1.11% colocalization of αPV/tdTomato, n = 3 mice; mean ± s.e.m.).

(C) Left: Representative confocal image of tdTomato fluorescence in a coronal section from a PV-tdTomato mouse containing BLA. Scale bar: 100 μm. Middle: Representative confocal images of colocalization experiment in a PV-tdTomato mouse. Left panel, tdTomato fluorescence. Middle panel, αSst or αPV immunofluorescence. Right panel, merge. Scale bar: 50 μm.

(D) Percent colocalization of αSst or αPV immunofluorescence with tdTomato fluorescence in PV-tdTomato mice (18.34 ± 2.24% colocalization of αSst/tdTomato, 90.82 ± 1.34% colocalization of αPV/tdTomato, n = 3 mice).

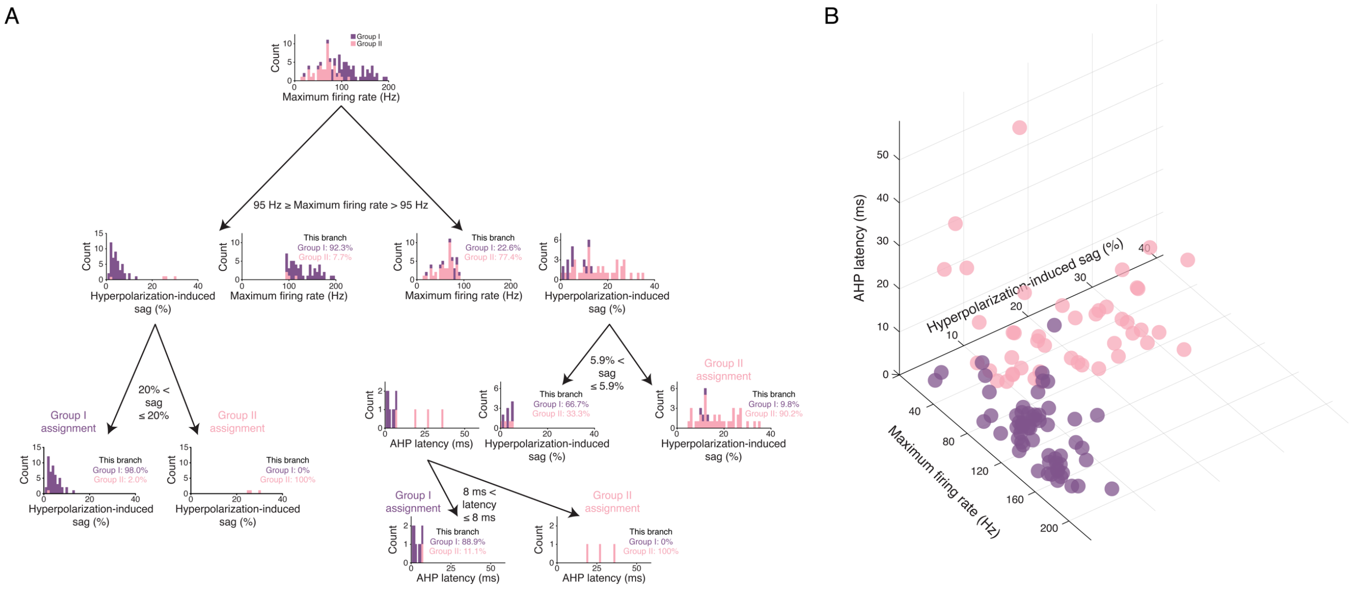


**Figure S3. Decision tree analysis to determine most salient parameters that discriminate Group I and II Sst⁺ INs. Related to Figure 3.**

(A) Results of the decision tree model. The model returned maximum firing rate, hyperpolarization-induced sag, and AHP latency as the most salient parameters for separating Group I and Group II $Sst^+$ INs. The tree shows the cut offs used to split the $Sst^+$ INs. The histograms show the counts of the two $Sst^+$ clusters at each branch of the decision tree (Group I: purple; Group II: pink).

(B) Scatter plot showing Group I (purple) and Group II (pink) $Sst^+$ IN maximum firing rates, hyperpolarization-induced sag, and AHP latency.

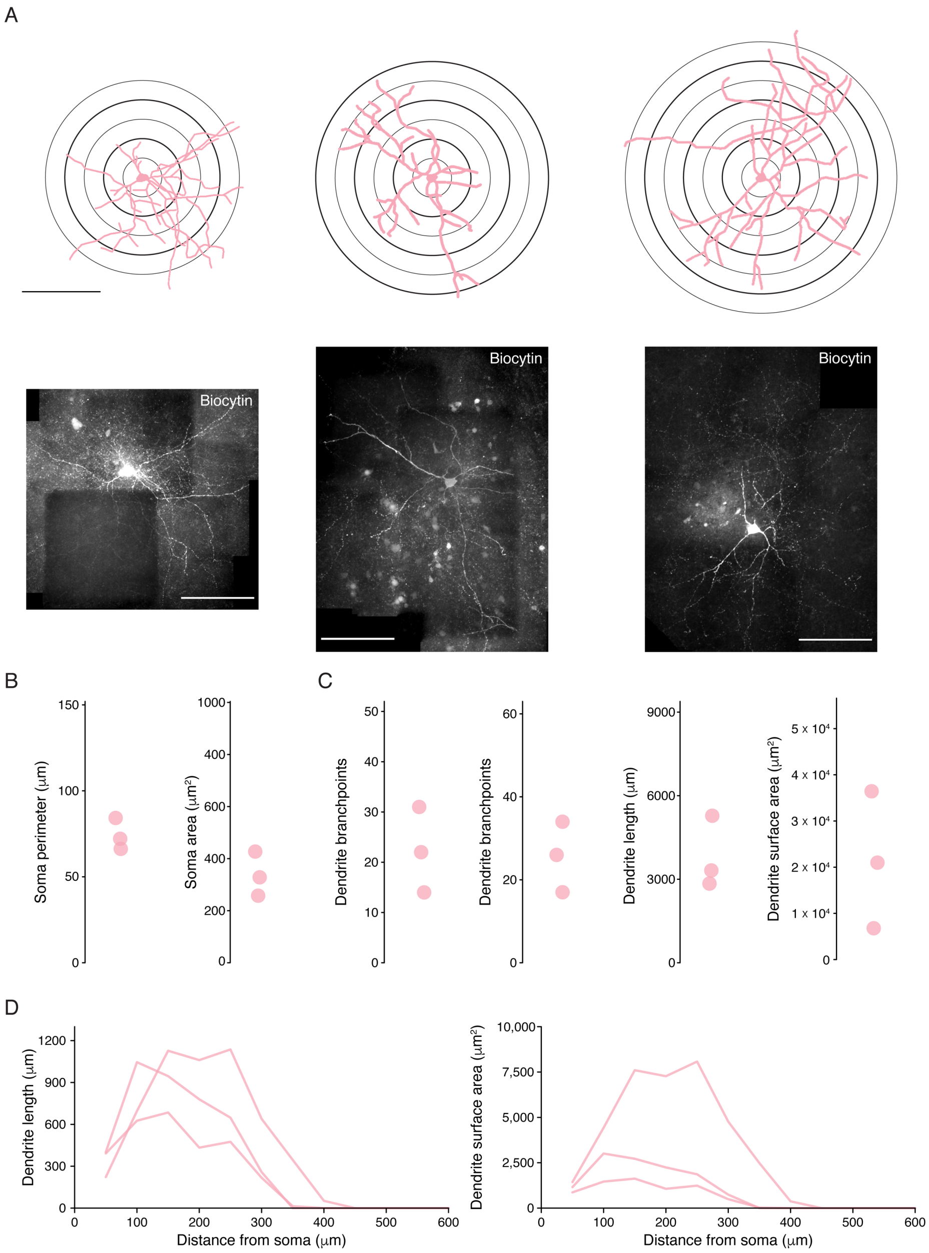


**Figure S4. Group II Sst⁺ IN membrane properties. Related to Figure 3.**

(A) Top: reconstructed soma and dendrites of all recovered Group II Sst⁺ INs (scale bar: 200 μm). Sholl rings shown beneath reconstruction. Bottom: confocal images of the corresponding biocytin filled neurons (scale bar: 200 μm).

(B) Soma morphology data for all recovered Group II Sst⁺ INs.

(C) Dendrite morphology data for all recovered Group II Sst⁺ INs.

(D) Sholl analysis of dendrite length (left) and surface area (right) as a function of distance from the soma in all recovered Group II Sst INs.

n = 3, 2.

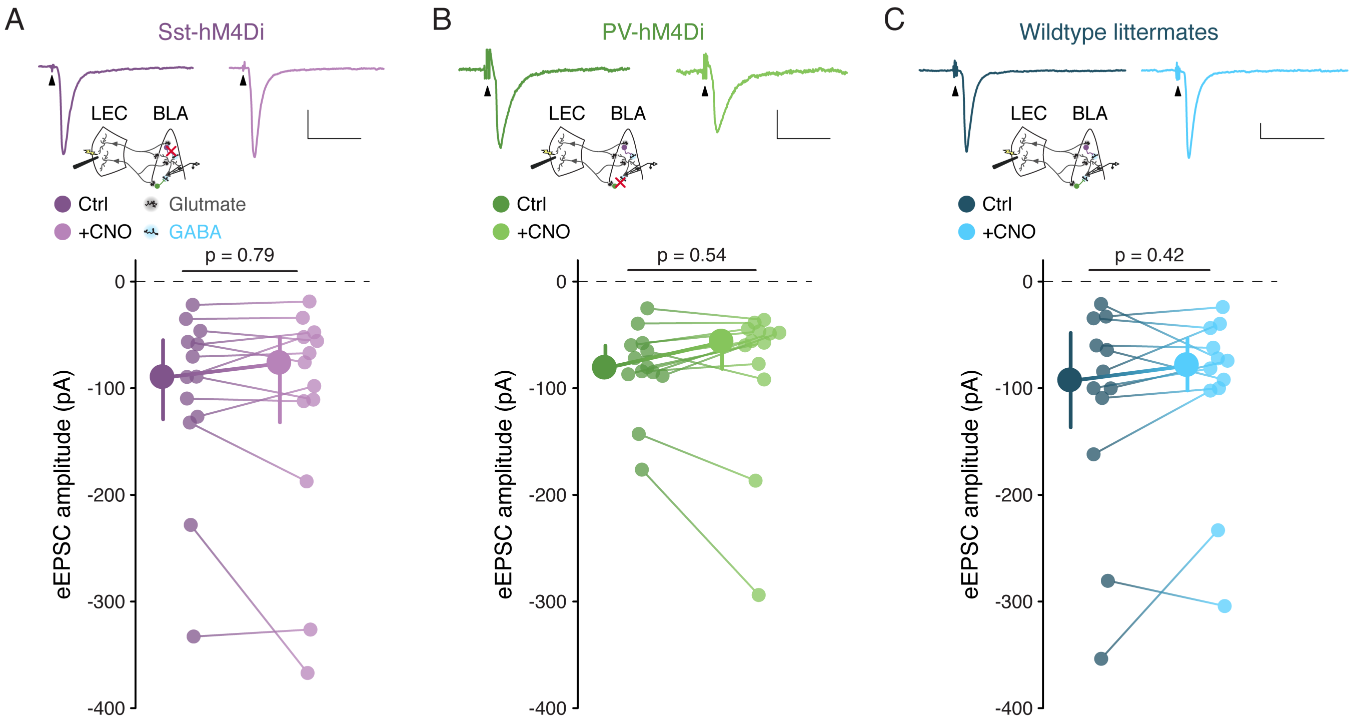


**Figure S5. Lack of an effect of 10 μM CNO on evoked EPSC amplitude. Related to Figure 5.**

(A) Top: Mean eEPSCs from a representative PN from a Sst-hM4Di mouse before and after bath application of CNO. Schematic of experiment below traces. Arrowheads denote stimulus artifacts (truncated). Scalebars: 100 pA, 50 ms. Bottom: $Sst^{+}$ IN inactivation has no effect on eEPSC amplitude ($eEPSC_{Ctrl}$ = -90.14 [-54.95/-129.09] pA, $eEPSC_{CNO}$ = -76.80 [-51.94/-131.96] pA, $p$ = 0.79, WSR test, n = 13 cells, 6 mice).

(B) As for (A), but in the PV-hM4Di mouse line. Scalebars: 20 pA, 50 ms. $PV^{+}$ IN inactivation has no effect on eEPSC amplitude ($eEPSC_{Ctrl}$ = -81.14 [-60.18/-88.34] pA, $eEPSC_{CNO}$ = -56.77 [-47.07/-81.82] pA, $p$ = 0.54 WSR test, n = 13 cells, 7 mice).

(C) As for (A), but in wildtype littermates of the Sst- and PV-hM4Di mice. Scalebars: 50 pA, 50 ms. 10 μM CNO has no effect on eEPSC amplitude in $hM4Di^{-/-}$ mice ($eEPSC_{Ctrl}$ = -93.18 [-48.19/-136.58] pA, $eEPSC_{CNO}$ = -78.85 [-53.80/-102.11] pA, $p$ = 0.42, WSR test, n = 12 cells, 6 mice).

Summary statistics presented in color as median with IQR.

| | Group I $Sst^+$ IN (n = 60) | | Group II $Sst^+$ IN (n = 45) | | $PV^+$ IN (n = 52) | | Statistical Comparisons |
|---|---|---|---|---|---|---|---|
| | *Mean/ Median* | *Variance* | *Mean/ Median* | *Variance* | *Mean/ Median* | *Variance* | |
| Membrane Resistance (MΩ) | 181.23[a,b] | 136.91/ 277.74 | 291.81[a,c] | 228.86/ 334.40 | 147.46[b,c] | 95.72/ 221.33 | KW: $p = 2.65 \times 10^{-10}$, $\alpha_{FDR}$: 0.05<br>[a]I vs II: $p = 3.18 \times 10^{-6}$<br>[b]I vs PV: $p = 0.011$<br>[c]II vs PV: $p = 2.29 \times 10^{-10}$ |
| Max. Firing Rate (Hz) | 121.14[a] | ± 4.87 | 66.75[a,b] | ± 3.18 | 116.24[b] | ± 7.29 | ANOVA: $p = 5.62 \times 10^{-11}$<br>[a]I vs II: $p = 9.70 \times 10^{-10}$<br>[b]II vs PV: $p = 4.22 \times 10^{-9}$ |
| Latency to First Action Potential (ms) | 84.65 | 57.40/ 175.70 | 85.10 | 60.75/ 133.30 | 131.15 | 70.40/ 229.10 | *NS*, KW: $p = 0.062$ |
| Action Potential Halfwidth (ms) | 0.50[a,b] | 0.43/0.59 | 0.70[a,c] | 0.60/0.87 | 0.44[b,c] | 0.40/0.50 | KW: $p = 4.34 \times 10^{-14}$, $\alpha_{FDR}$: 0.05<br>[a]I vs II: $p = 2.21 \times 10^{-9}$<br>[b]I vs PV: $p = 5.56 \times 10^{-4}$<br>[c]II vs PV: $p = 3.86 \times 10^{-12}$ |
| Firing Rate Adaptation | 0.69[a] | 0.59/0.89 | 0.53[a,b] | 0.31/0.63 | 0.74[b] | 0.37/0.92 | KW: $p = 1.89 \times 10^{-4}$, $\alpha_{FDR}$: 0.033<br>[a]I vs II: $p = 3.36 \times 10^{-5}$<br>[b]II vs PV: $p = 0.0024$ |
| Action Potential Broadening | 1.00 | 1.00/1.20 | 1.00 | 1.00/1.14 | 1.00 | 1.00/1.23 | *NS*, KW: $p = 0.34$ |
| Action Potential Threshold (mV) | -35.28 | -39.47/ -32.22 | -34.83 | -36.76/ -30.77 | -34.74 | -38.79/ -31.32 | *NS*, KW: $p = 0.51$ |
| Action Potential Amplitude (mV) | 46.80[a] | ± 1.11 | 56.34[a,b] | ± 1.91 | 50.09[b] | ± 1.46 | ANOVA: $p = 5.18 \times 10^{-5}$<br>[a]I vs II: $p = 1.47 \times 10^{-5}$<br>[b]II vs PV: $p = 0.012$ |
| Amplitude Adaptation | 0.88[a] | ± 0.014 | 0.82[a,b] | ± 0.018 | 0.89[b] | ± 0.017 | ANOVA: $p = 0.011$<br>[a]I vs II: $p = 0.038$<br>[b]II vs PV: $p = 0.011$ |
| AHP magnitude (mV) | 20.04 | ± 0.52 | 19.42 | ± 0.90 | 21.20 | ± 0.58 | *NS*, ANOVA: $p = 0.17$ |
| ΔAHP (mV) | 2.72 | 1.69/4.05 | 2.84[a] | 1.85/5.06 | 2.06[a] | 1.08/3.22 | KW: $p = 0.018$, $\alpha_{FDR}$: 0.017<br>[a]II vs PV: $p = 0.011$ |
| AHP Latency (ms) | 3.20[a] | 2.43/4.49 | 6.80[a,b] | 2.83/15.58 | 3.00[b] | 2.00/3.64 | KW: $p = 2.52 \times 10^{-5}$, $\alpha_{FDR}$: 0.033<br>[a]I vs II: $p = 8.24 \times 10^{-4}$<br>[b]II vs PV: $p = 1.79 \times 10^{-5}$ |
| Hyperpolarization-induced Sag (%) | 4.41[a] | 3.07/6.89 | 16.45[a,b] | 10.24/ 25.17 | 5.38[b] | 3.80/7.97 | KW: $p = 1.68 \times 10^{-12}$, $\alpha_{FDR}$: 0.033<br>[a]I vs II: $p = 2.74 \times 10^{-11}$<br>[b]II vs PV: $p = 1.00 \times 10^{-9}$ |
| Rebound Action Potentials | 0.00[a] | 0.00/0.00 | 0.00[a,b] | 0.00/1.25 | 0.00[b] | 0.00/0.00 | KW: $p = 1.92 \times 10^{-9}$, $\alpha_{FDR}$: 0.033<br>[a]I vs II: $p = 2.15 \times 10^{-7}$<br>[b]II vs PV: $p = 5.68 \times 10^{-6}$ |
| $\tau_{Membrane}$ (ms) | 11.74[a] | 9.57/16.05 | 19.53[a,b] | 16.36/ 26.18 | 11.25[b] | 9.74/ 14.26 | KW: $p = 2.70 \times 10^{-11}$, $\alpha_{FDR}$: 0.033<br>[a]I vs II: $p = 6.43 \times 10^{-9}$<br>[b]II vs PV: $p = 1.18 \times 10^{-10}$ |

**Table S1. Differences in active and passive membrane properties among BLA $Sst^+$ and $PV^+$ INs. Related to figure 3.**

Normal data are presented as mean ± s.e.m. with differences tested using a 1-way ANOVA with Tukey's honestly significant difference post hoc test. Non-normal data are presented as median and IQR with differences tested using a KW test with MWU tests between groups as a post hoc. Multiple comparisons with the MWU tests were corrected using the false discovery rate method where $p < \alpha_{FDR}$ was considered significant.
