## Supplementary material for "Cell type specific control of basolateral amygdala plasticity via entorhinal cortex-driven feedforward inhibition": Key Resources Table

| REAGENT or RESOURCE | | SOURCE | | IDENTIFIER |
| --- | --- | --- | --- | --- |
| Antibodies | | | | |
| Rat anti-Sst | | Millipore | | Cat#MAB354; RRID: AB_2255365 |
| Guinea pig anti-PV | | Synaptic Systems | | Cat#195004; RRID: AB_2156476 |
| Donkey anti-rat Alexa Fluor 488 | | Jackson ImmunoResearch | | Cat#712-545-150 |
| Donkey anti-guinea pig Alexa Fluor 488 | | Jackson ImmunoResearch | | Cat#706-545-148 |
| Streptavidin Conjugated Alexa Fluor 488 | | ThermoFisher Scientific | | Cat#S11223 |
| Chemicals, Peptides, and Recombinant Proteins | | | | |
| iFX-enhancer | | Invitrogen | | Cat#I36933 |
| Picric acid | | Electron Microscopy Sciences | | Cat#I9556 |
| Glutaraldehyde | | Sigma | | Cat#G-7651 |
| Sucrose | | Sigma-Aldrich | | Cat#S7903 |
| Glucose | | Sigma-Aldrich | | Cat#G8270 |
| Sodium chloride | | Sigma-Aldrich | | Cat#S9888 |
| Potassium chloride | | Sigma-Aldrich | | Cat#P9333 |
| Sodium phosphate monobasic dihydrate | | Sigma-Aldrich | | Cat#71505 |
| Sodium bicarbonate | | Sigma-Aldrich | | Cat#S6297 |
| Calcium chloride | | Sigma-Aldrich | | Cat#21115 |
| Magnesium chloride | | Sigma-Aldrich | | Cat#68475 |
| Cesium methanesulfonate | | Acros Organics | | CAS: 2550-61-0 |
| HEPES | | Sigma-Aldrich | | Cat#H3375 |
| EGTA | | Sigma-Aldrich | | Cat#E3889 |
| Sodium phosphocreatine | | Sigma-Aldrich | | Cat#P7936 |
| QX-314 (Lidocaine N-ethyl bromide) | | Sigma-Aldrich | | Cat#L5783 |
| Magnesium-ATP | | Sigma-Aldrich | | Cat#A9187 |
| Sodium-GTP | | Sigma-Aldrich | | Cat#G8877 |
| Potassium gluconate | | Sigma-Aldrich | | Cat#G4500 |
| DNQX | | Tocris | | Cat#0189; CAS: 2379-57-9 |
| D-APV | | Tocris | | Cat#0106; CAS: 79055-68-8 |
| Gabazine (SR 95531 hydrobromide) | | Tocris | | Cat#1262; CAS: 104104-50-9 |
| Clozapine-*N*-oxide | | Enzo Life Sciences | | Cat#BML-NS105 |
| Phosphate Buffered Saline (PBS) Tablets, 100 mL | | VWR | | Cat#E404-200TABS |
| Triton X-100 | | VWR | | Cat#0694-1L |
| Normal Donkey Serum | | Jackson Immuno-research Labs Inc. | | Cat#017-000-121 |
| Bovine Serum Albumin | | Sigma-Aldrich | | Cat#A9647-100G |
| Prolong Gold antifade reagent | | Invitrogen | | Cat#P36934 |
| Nail Polish | | Electron Microscope Sciences | | Cat#72180 |
| Biocytin | | ThermoFisher | | Cat#28022 |
| Deposited Data | | | | |
| Raw and analyzed data | | This paper | | Available on request |
| Experimental Models: Organisms/Strains | | | | |
| Mouse: C57BL/6J | Jackson Lab | | RRID: IMSR_JAX:000664 | |
| Mouse: Sst^tm2.1(cre)Zjh^/J | Jackson Lab | | RRID: IMSR_JAX:013044 | |
| Mouse: B6;129P2-Pvalb^tm1(cre)Arbr^/J | Jackson Lab | | RRID: IMSR_JAX:008069 | |
| Mouse: B6;129S6-Gt(ROSA)26Sor^tm9(CAG-tdTomato)Hze^/J | Jackson Lab | | RRID: IMSR_JAX:007905 | |
| Mouse: B6N.129-Gt(ROSA)26Sor^tm1(CAG-CHRM4*,-mCitrine)Ute^/J | Jackson Lab | | RRID: IMSR_JAX:026219 | |
| Software and Algorithms | | | | |
| pClamp 10.6 | | Molecular Devices | | RRID: SCR_011323 |
| MATLAB_R2018a | | Mathworks | | RRID: SCR_001622 |
| R | | <http://www.r-project.org/> | | RRID: SCR_001905 |
| Illustrator | | Adobe | | RRID: SCR_010279 |
| Photoshop | | Adobe | | RRID: SCR_014199 |
| SlideBook 6.0 | | Intelligent Imaging Innovations (3i) | | RRID: SCR_014300 |
| Fiji | | Fiji | | RRID: SCR_002285 |
| Grid/Collection Stitching | | Fiji | | RRID: SCR_016568 |
| Neurolucida | | MBF Biosciences | | RRID: SCR_001775 |
